# The evolution of two transmissible cancers in Tasmanian devils

**DOI:** 10.1101/2022.05.27.493404

**Authors:** Maximilian R. Stammnitz, Kevin Gori, Young Mi Kwon, Ed Harry, Fergal J. Martin, Konstantinos Billis, Yuanyuan Cheng, Adrian Baez-Ortega, William Chow, Sebastien Comte, Hannes Eggertsson, Samantha Fox, Rodrigo Hamede, Menna E. Jones, Billie Lazenby, Sarah Peck, Ruth Pye, Michael A. Quail, Kate Swift, Jinhong Wang, Jonathan Wood, Kerstin Howe, Michael R. Stratton, Zemin Ning, Elizabeth P. Murchison

## Abstract

Tasmanian devils have spawned two transmissible cancer lineages, named devil facial tumour 1 (DFT1) and devil facial tumour 2 (DFT2). We investigated the genetic diversity and evolution of these clones by analysing 78 DFT1 and 41 DFT2 genomes relative to a newly assembled chromosome-level reference. Time-resolved phylogenetic trees reveal that DFT1 first emerged in 1986 (1982-1989), and DFT2 in 2011 (2009-2012). Subclone analysis documents transmission of heterogeneous cell populations. DFT2 has faster mutation rates than DFT1 across all variant classes, including substitutions, indels, rearrangements, transposable element insertions and copy number alterations, and we identify a hypermutated DFT1 lineage with defective DNA mismatch repair. Several loci show plausible evidence of positive selection in DFT1 or DFT2, including loss of chromosome Y and inactivation of *MGA*, but none are common to both cancers. This study illuminates the parallel long-term evolution of two transmissible cancers inhabiting a common niche in Tasmanian devils.

---

Transmissible cancers are contagious somatic cell lineages that spread through populations by the physical transfer of living cancer cells. Although few such diseases are known in nature, Tasmanian devils (*Sarcophilus harrisii*), marsupial carnivores endemic to the Australian island of Tasmania, host at least two transmissible cancer clones. These cancers, known as devil facial tumour 1 (DFT1) and devil facial tumour 2 (DFT2), both primarily cause malignant facial and oral tumours that are spread by biting (Figure 1A) (*1–3*). DFT1 was first observed in 1996 in north-eastern Tasmania and has subsequently spread widely (*4, 5*); DFT2, on the other hand, was discovered in 2014 on the D’Entrecasteaux Channel Peninsula in Tasmania’s south-east, and is believed to remain confined to this area (*3, 6, 7*). Both DFT1 and DFT2 are usually fatal, and rapid Tasmanian devil population declines associated with DFT1 have led to concern for conservation of the species (*4, 5, 8*).

**Fig. 1:**
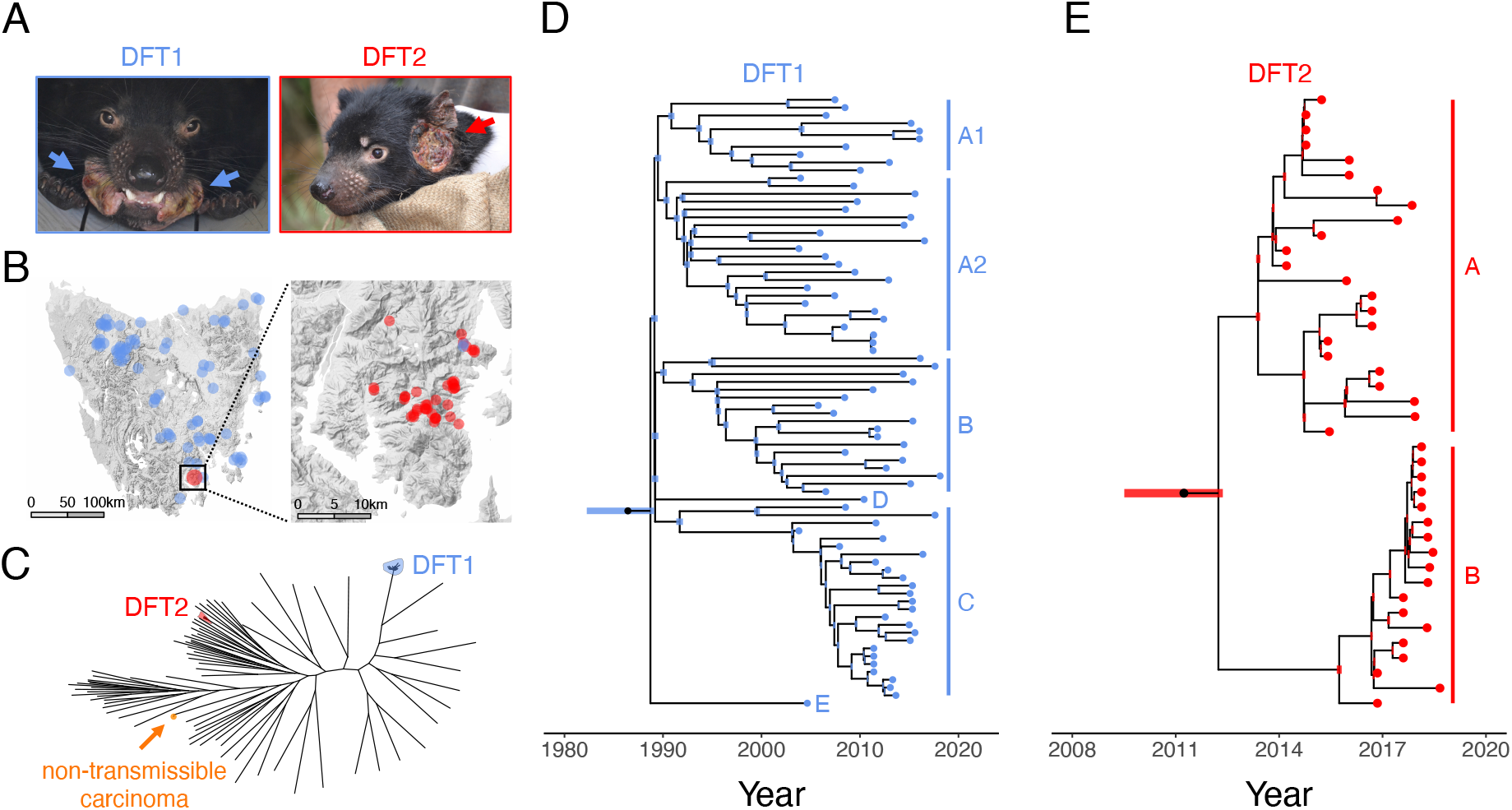
DFT1 and DFT2 phylogenies. (**A**) Representative photographs of animals infected with DFT1 and DFT2. (**B**) Sampling locations of 78 DFT1 and 41 DFT2 tumours included in the study. (**C**) Maximum likelihood phylogenetic tree constructed using 104,799 somatic and 1,070,436 germline substitutions from 38 DFT1s, 12 DFT2s, a single non-transmissible carcinoma and 79 Tasmanian devils; only the subset of DFT1s and DFT2s with tumour purity ≥75% were included. Black unlabelled tips represent Tasmanian devils and shaded tips represent those belonging to DFT1, DFT2 or the non-transmissible carcinoma. Branch lengths are uninformative. High resolution labelled tree available in Figure S2. (**D** and **E**) Time-resolved phylogenetic trees for DFT1 and DFT2 constructed using 171,283 and 21,252 somatic substitution mutations, respectively. Tumour clades are labelled (A1, A2, B, C, D, E in DFT1; A, B in DFT2). Bars at internal nodes represent 95% Bayesian credible intervals around date estimates. Bars at root nodes represent 95% Bayesian credible intervals around date estimates, incorporating uncertainty in somatic/germline assignment of substitutions shared by all tumours within a clone and absent from all normal Tasmanian devils. Dating is based on tumour sampling dates and does not account for the pretransmission interval, the offset between date of clone emergence and date of sampling; this is of relevance because bulk tissue sequencing captures only clonal mutations or those present in sizeable subclones (*51*). High resolution labelled trees available in Figures S3 and S5.

The emergence of two transmissible cancers in Tasmanian devils suggests that the species is particularly susceptible to this type of disease. Indeed, DFT1 and DFT2 appear to be independent occurrences of the same pathological process, and their comparison may illuminate the constraints of the biological niche that they inhabit. DFT1 and DFT2 are both undifferentiated Schwann cell cancers with similar dependence on receptor tyrosine kinase signalling (*9–12*). DFT1 first arose from the cells of a female “founder devil” and equally affects male and female devil hosts (*2, 13–15*); DFT2, on the other hand, originated from a male devil and shows preference for male hosts, perhaps due to immunogenicity of chromosome Y-derived antigens in female hosts (*3, 7, 10*). Both cancers escape the allogeneic immune system, and, in DFT1, this is mediated by transcriptional repression of major histocompatibility complex (MHC) class I genes (*16*); in DFT2, however, cell surface MHC class I molecules are usually detectable, and high similarity between expressed tumour and host MHC class I alleles may underlie the lack of immune rejection (*17*). The genomes of DFT1 and DFT2 show comparable mutational patterns, but no common positively selected “driver” mutations have been detected (*10*). Furthermore, whereas DFT1 has split into several spatially defined sublineages during its spread through Tasmania (*18*), little is known about the clonal diversity of DFT2.

In addition to their importance as threats to animal health and their intrinsic interest as unusual pathogens, transmissible cancers provide an opportunity to study how mutations in cancer accumulate with time. Most human cancer studies involve the analysis of tumour biopsies collected either at a single session, or at time-points separated by short intervals. The long-term survival of DFT1 and DFT2 permits repeated sampling of the same cancer lineages through decades, enabling direct investigation of variation in mutation rates, together with those of their constitutive mutational signatures, within and between clones.

Here, we describe high-coverage whole genome sequences of 78 DFT1 and 41 DFT2 tumours, as well as that of a single non-transmissible carcinoma and a panel of 80 normal Tasmanian devil genomes, analysed relative to a newly assembled, highly contiguous Tasmanian devil reference genome. By capturing the somatic genetic diversity present within the DFT1 and DFT2 lineages, our goal was to understand the dynamics of these diseases’ emergence and spread, to estimate their mutation rates, and to characterise their long-term patterns of evolution. By intersecting findings from different Tasmanian devil cancers, we identify genomic events that underpin transmissible cancer in this species. Our analysis provides detailed insight into the evolution and diversification of two parallel cancer clones that have survived in a transmissible niche.

## Results

### A new reference genome for the Tasmanian devil

The previous Tasmanian devil reference genome, DEVIL7.0, was highly fragmented (35,534 chromosomal scaffolds) (*13*). In order to produce an improved genome assembly for the species, we extracted high molecular weight DNA from the female fibroblast cell line that was previously used in DEVIL7.0. We sequenced this to 76-fold and 12-fold coverage using long-read (fragment N50: 9.05 kilobases, kb) and ultra-long read (N50: 57.13 kb) sequencing technology (*19*). In addition, DNA was analysed using optical mapping, linked-read sequencing and high dimension conformation capture (Hi-C). A new reference genome assembly, mSarHar1.11, was generated by combining these data (Table 1, Table S1, Figure S1). Notably, 99.8 percent of bases were placed on one of seven scaffolds, corresponding to the six devil autosomes and chromosome X.

**Table 1:** mSarHar1.11 Tasmanian devil reference genome assembly and annotation metrics. mSarHar1.11 is compared with the previous reference assembly, DEVIL7.0 (*13*). Mb, megabase.

|  | DEVIL7.0 | mSarHar1.11 |
| --- | --- | --- |
| Contigs (N50) | 237,291 (0.02 Mb) | 445 (63.34 Mb) |
| Largest contig | 0.19 Mb | 195.48 Mb |
| Chromosomal scaffolds (N50) | 35,534 (1.85 Mb) | 7 (611.35 Mb) |
| Largest chromosomal scaffold | 5.32 Mb | 716.41 Mb |
| Repeat-masked genome | 39.6% | 44.3% |
| Coding genes | 18,788 | 19,228 |
| Non-coding genes | 1,490 | 4,336 |

Genome annotation was performed using the Ensembl gene annotation pipeline (*20*) guided by a newly sequenced Tasmanian devil multi-tissue transcriptome atlas, yielding 19,228 protein-coding gene models (Table S1).

### DFT1 and DFT2 phylogenies

In order to investigate genetic variation within Tasmanian devil transmissible cancers, we sequenced the whole genomes of 63 DFT1s and 39 DFT2s (Figure 1A) to a median depth of 83x, and analysed these alongside 15 DFT1 and 2 DFT2 publicly available genomes (Table S2). The DFT1s were primarily selected to capture genetic and spatiotemporal diversity in this clone (Figure 1B, Table S2). These included representatives of the six major clades (A1, A2, B, C, D and E) (*18*) and were collected from 38 locations between 2003 and 2018. For DFT2, on the other hand, we sequenced all available tumours sampled between 2014 and 2018, all occurring within DFT2’s known range on the D’Entrecasteaux Channel Peninsula (Figure 1B). Some subsets of DFT1 and DFT2 tumours were derived from the same individual hosts, including sets of matched primary facial tumours and internal metastases, as well as samples from distinct facial or body tumours occurring in single hosts (Table S2). In addition, we sequenced a non-transmissible anal sac carcinoma sampled from a captive Tasmanian devil, and analysed genomes from 80 normal Tasmanian devils including matched hosts (71 newly sequenced, 9 publicly available; Table S2).

Single-base substitutions were called in each sample, and normal Tasmanian devil genomes were used to identify and exclude germline substitutions from tumour sequences. This yielded 205,890, 23,152 and 5,764 somatic substitutions in DFT1, DFT2 and in the non-transmissible anal sac carcinoma respectively, as well as 1,458,776 germline variants (Table S3). Analysis of the latter revealed a median of 0.132 heterozygous sites per kilobase (range 0.083-0.153) in the sampled population of Tasmanian devils, with the DFT1 and DFT2 founder devils both falling within this range (Table S3).

We confirmed the independent clonal origins of DFT1 and DFT2 by constructing a maximum likelihood tree using substitutions from both tumour and normal samples. As expected, DFT1 and DFT2 tumours each clustered into distinct groups whose positions relative to normal animals are consistent with the notion that these clones’ founder devils originated in north-eastern Tasmania (DFT1) or on the D’Entrecasteaux Channel Peninsula (DFT2) (Figure 1C, Figure S2) (*3, 4, 10*).

Time-resolved phylogenetic trees were generated for DFT1 and DFT2 with substitution mutation rates inferred using tumour sampling dates (Figure 1D and 1E). Assuming a constant mutation rate, DFT1 was estimated to have arisen in 1986 (95% Bayesian credible interval 1982-1989), implying a substantial delay from its emergence until its first observation in 1996 (Figure 1D, Figure S3) (*4*). The DFT1 tree showed the expected arrangement of the six identified tumour clades (*18*), and revealed that these split from one another very early in DFT1 evolution in a rapid diversification event that almost certainly involved a single tumour donor (Figure S4). DFT2, on the other hand, is estimated to have first emerged in 2011 (95% Bayesian credible interval 2009-2012). It subsequently split into two major sympatric groups which we term DFT2 clades A and B (Figure 1E, Figure S5). The potential for individual devils to be coinfected with distinct lineages of DFT1 (*18*), DFT2, or both (*10*) is apparent. The presence of true- or near-polytomies evident on both the DFT1 and DFT2 phylogenetic trees, defined by very short internal branches (Figure 1D and 1E), suggests that it may not be uncommon for infectious devils to transmit their tumour to more than two secondary hosts. Such events may, however, be enriched at early time-points in the trees due to survivorship bias (*21*).

### Intra-tumour genetic heterogeneity in DFT1 and DFT2

Bulk sequencing of tumour tissue, as performed here, will capture only clonal mutations or those present in sizeable subclones. Where present, however, the distribution of subclones among tumours could be informative about the clonality of transmission in DFT1 and DFT2.

We screened tumours for subclones by searching for mutation populations showing unexpected allele fractions. One DFT2 tumour, 1509T1, was found to be composed of two subclonal cell populations represented at roughly 60% and 40% frequency, respectively. We computationally isolated these subclones, and inspection of their positions on the DFT2 phylogenetic tree revealed that they belonged to separate DFT2 clade B sublineages, which we term DFT2-B2 and DFT2-B3 (Figure 2A and 2B). Indeed, mutations defining each subclone were observed clonally in related contemporaneous tumours from different hosts. These data are compatible with a model whereby an earlier donor tumour contained cells belonging to both DFT2-B2 and DFT2-B3; onward transmission founded descendent tumours composed of either DFT2-B2 or DFT2-B3 cells, or, in the case of 1509T1, a mixture of both DFT2-B2 and DFT2-B3 cells (Figure 2C).

**Fig. 2:**
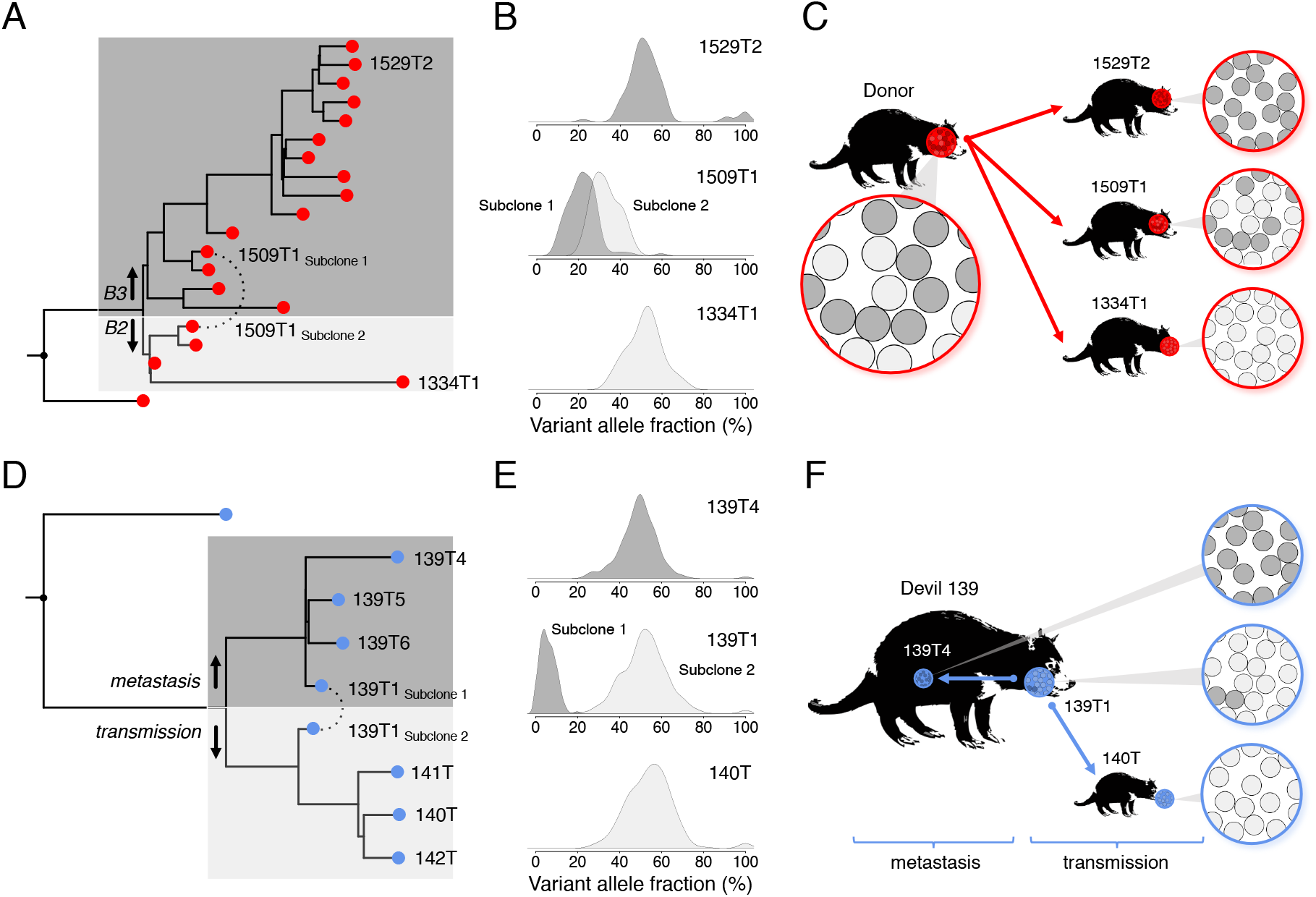
Intra-tumour genetic heterogeneity in DFT1 and DFT2. (**A-C**) An example of heterogeneous cell transmission in DFT2. 1509T1 is a DFT2 clade B tumour composed of two detectable subclonal cell populations, 1509T1_Subclone1_ and 1509T1_Subclone2_. (**A**) Computational separation of 1509T1_Subclone1_ and 1509T1_Subclone2_ and inclusion on a phylogenetic tree revealed subclone membership of distinct clade subgroups, DFT2-B3 and DFT2-B2. Branch lengths are proportional to number of substitution variants. (**B**) Variant allele distribution of 1509T1, together with those of representative DFT2-B3 (1529T2) and DFT2-B2 (1334T1) tumours; only variants occurring after the split between DFT2-B3 (dark grey) and DFT2-B2 (light grey) are included. As tumours are diploid, most mutations occur in the heterozygous state and would be expected to be found at 50% proportion. (**C**) Model illustrating transmission of DFT2 from an earlier donor devil, which carried both DFT2-B3 (dark grey) and DFT2-B2 (light grey) cells, to recipient devils. Recipient tumours are composed either of clonal populations of DFT2-B3 (upper, dark grey, 1529T2), clonal populations of DFT2-B2 (lower, light grey, 1334T1) or a subclonal mixture of DFT2-B3 and DFT2-B2 (middle, mixture of light grey and dark grey, 1509T1). Arrows do not necessarily represent direct transmission. (**D**-**F**) An example of differential transmission and metastasis of subclones in DFT1. 139T1 is a DFT1 facial tumour composed of two detectable subclonal cell populations, 139T1_Subclone1_ and 139T1_Subclone2_, represented by dark grey and light grey shading, respectively. (**D**) 139T1_Subclone1_ clusters phylogenetically with a set of internal metastases sampled from the same individual (139T4, 139T5, 139T6), and 139T1_Subclone2_ clusters with facial tumours sampled from three devils (140T, 141T, 142T) involved in a DFT transmission chain (Figure S6). Branch lengths are proportional to number of substitution variants. (**E**) Variant allele distribution of 139T1, together with those of a representative metastasis involving the same host (139T4), and a representative tumour secondary to transmission involving a different host (140T); only variants occurring after the split between the metastases (dark grey) and transmission (light grey) are included. As tumours are diploid, most mutations occur in the heterozygous state and would be expected to be found at 50% proportion. (**F**) Model illustrating differential spread of subclones. 139T1_Subclone1_ (dark grey) and 139T1_Subclone2_ (light grey) are both represented in tumour 139T1. Cells belonging to 139T1_Subclone1_ seeded internal metastases (represented by 139T4), whereas cells from 139T1_Subclone2_ were transmitted onwards to recipient devils (represented by 140T). Further details available in Figure S6.

We similarly investigated intratumour heterogeneity in DFT1 using a closely related set of tumours that were part of a series of direct transmission events (Figure 2D, Figure S6). This case involved a female devil with a facial tumour and several metastases. Cells were transmitted from this female’s facial tumour to her unweaned male offspring, which, once weaned, further transmitted his tumour to two additional hosts while the group was housed together in captivity (Figure S6). The index female’s facial tumour was composed of two detectable subclones at roughly 90% and 10% proportions which clustered with the tumour of the offspring and with her metastases, respectively (Figure 2D and 2E). This suggests that two distinct cell lineages, both represented within the index facial tumour, differentially contributed to metastatic dissemination and onward transmission (Figure 2F).

These case studies hint at the genetic heterogeneity present within individual DFT tumours, and, in the DFT2 example, imply that this diversity can be maintained across transmission bottlenecks. Thus, at least in some cases, DFT tumours are seeded by more than one cell.

### DFT1 and DFT2 substitutions and indels

To obtain an overview of the mutational processes operating in Tasmanian devil cancers, we inspected each tumour’s mutational spectrum, a representation of the distribution of mutations across the six base substitution classes, displayed together with their immediate 5’ and 3’ base contexts. Such spectra can be decomposed into their constituent mutational signatures, patterns of co-occurring mutation types which reflect the activities of underlying endogenous or exogenous mutational processes (*22*). As expected, DFT1 and DFT2, as well as the single non-transmissible anal sac carcinoma, showed evidence for the presence of two known mutational signatures, single base substitution signatures 1 (SBS1) and 5 (SBS5), which are found almost universally in human cancer (*23*), and have been described previously in Tasmanian devil tumours (Figure 3A, Figure S7) (*10*). SBS1 is characterised by C>T mutations at <u>C</u>pG dinucleotide contexts (mutated base underlined) and is believed to primarily arise due to spontaneous deamination of 5’-methylcytosine (*22*). SBS5, on the other hand, shows little base specificity and its aetiology is poorly understood (*23, 24*). Consistent with a previous report (*10*), no evidence of ultraviolet light mutagenesis was detectable in DFT1 or DFT2 mutation patterns, indicating that the cells that transmit DFT are not usually exposed to sunlight. Patterns of short insertions and deletions (indels) in DFT1 and DFT2 revealed imprints of Indel signatures 1 (ID1) and 2 (ID2) in both cancers, although ID1 dominated in DFT1 (66% ID1, 34% ID2) whereas ID1 and ID2 were present at similar proportions in DFT2 (47% ID1, 53% ID2; Figure 3B, Figure S8). These signatures are defined by the accumulation of insertions (ID1) or deletions (ID2) of single thymine or adenine bases occurring at mononucleotide tracts, and arise through polymerase slippage involving the nascent (ID1) or the template (ID2) DNA strand (*23*).

**Fig. 3:**
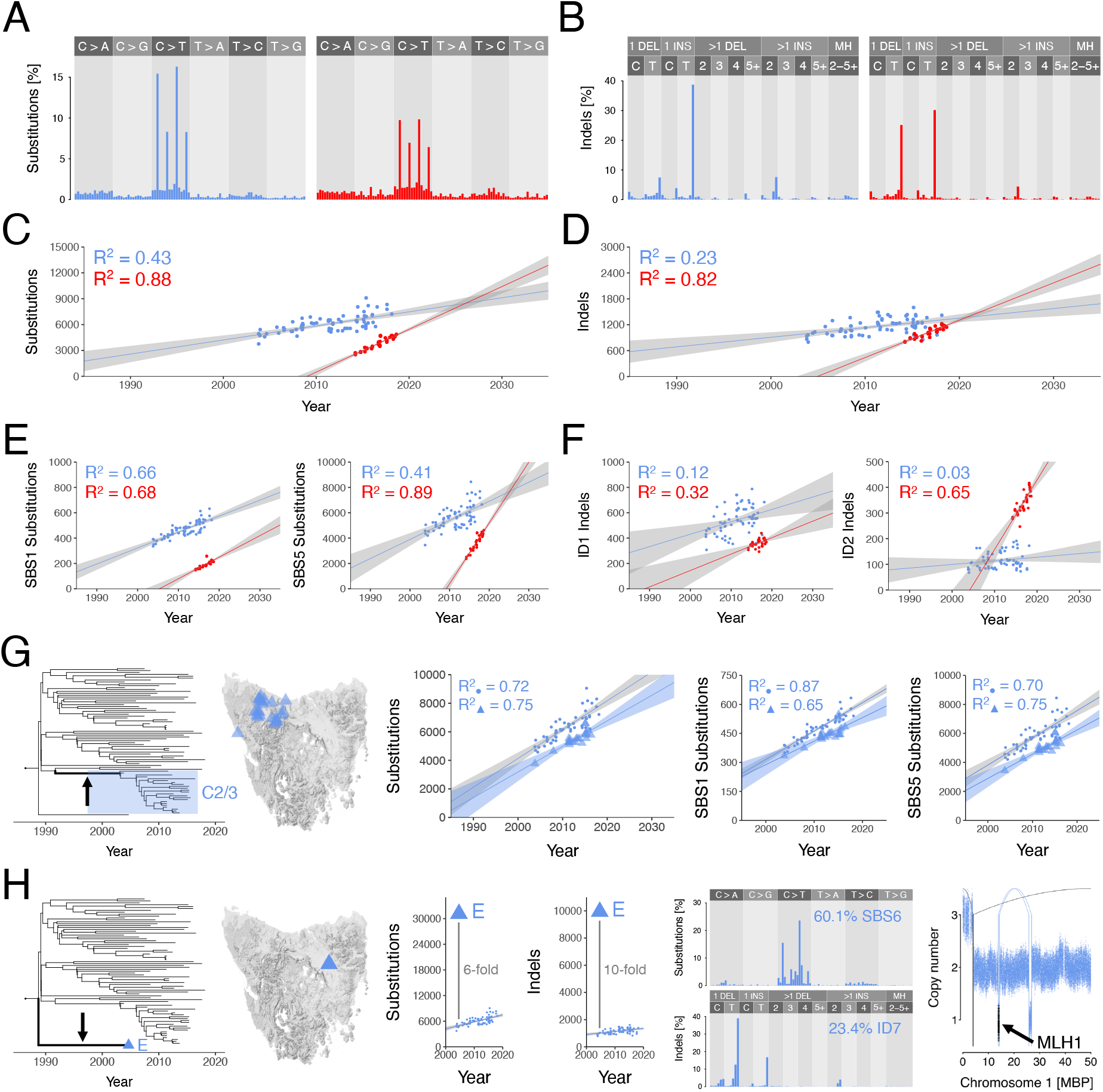
DFT1 and DFT2 substitutions and indels. (**A** to **B**) Mutational spectra for somatic substitutions (A) and indels (B) in DFT1 (blue, N = 176,428 substitutions, N = 22,479 indels; variants unique to DFT1 clade E were excluded) and DFT2 (red, N = 23,152 substitutions, N = 4,054 indels). Fully labelled plots available in Figures S7 and S8. (**C** to **D**) Rate of accumulation of substitutions (C) and indels (D) in DFT1 excluding clade E (blue) and DFT2 (red). Each point represents a tumour, plotted by sampling date. Lines represent linear regression, grey shading 95% confidence interval. (**E**) Rate of accumulation of substitution mutations corresponding to mutational signatures SBS1 (left) and SBS5 (right) in DFT1 excluding clade E (blue) and DFT2 (red). Each point represents a tumour, plotted by sampling date. Lines represent linear regression, grey shading 95% confidence interval. (**F**) Rate of accumulation of substitution mutations corresponding to mutational signatures ID1 (left) and ID2 (right) in DFT1 excluding clade E (blue) and DFT2 (red). Each point represents a tumour, plotted by sampling date. Lines represent linear regression, grey shading 95% confidence interval. (**G**) Transient reduction in DFT1 substitution mutation rate occurring within phylogenetic branch leading to DFT1 clade C2/3 (arrow and shading, left); tumours in DFT1 clade C2/3 occur in Tasmania’s north-west (map). Overall mutation rate reduction (centre) is attributable to both mutational signatures SBS1 and SBS5 (second from right, right); each point represents a tumour, plotted by sampling date, with clade C2/3 tumours represented as triangles. Lines represent linear regression, grey shading 95% confidence interval. (**H**) The single representative of DFT1 clade E, sampled in north-east Tasmania (tree, map) has elevated numbers of substitution and indel mutations; central plots show numbers of substitutions (left) and indels (right) in all DFT1 tumours plotted by sampling date, with clade E tumour represented by triangle. Clade E has distinctive substitution and indel mutational spectra, with at least 60% of the spectrum explained by signature SBS6 (second from right; fully labelled plots available in Figure S9). Clade E carries a deletion encompassing the *MLH1* locus (right; dots represent normalised read coverage within 1 kilobase genomic windows, with windows including *MLH1* shaded in black; MBP, mega base pairs; connecting arcs represent rearrangements). High-resolution images and source data available in Figures S7-S9, and Table S3.

Mutational signatures SBS1, SBS5, ID1 and ID2 all present “clock-like” properties in human cells, showing linear correlation with donor age (*23, 25, 26*). Their rates vary widely among tissues, and, whereas the rates of SBS1, ID1 and ID2 correlate with one another and are believed to reflect the number of mitoses that a cell has experienced, SBS5 rate is independent of these (*23*). We characterised overall substitution and indel rates, as well as rates of SBS1, SBS5, ID1 and ID2 in DFT1 and DFT2 by regressing the number of mutations attributable to each signature in each tumour against sampling date (Figure 3C-3F). These analyses revealed that overall substitution and indel mutation rates in DFT2 were 3.0 and 3.9 times higher, respectively, than those of DFT1 (Table 2). The magnitude of these differences was, however, signature-specific. SBS1 and ID1 accumulate only moderately faster in DFT2 than in DFT1, but rates of SBS5 and ID2 are both considerably higher in DFT2 than in DFT1 (Table 2, Figure 3E and 3F, Table S3).

**Table 2:** Summary of DFT1 and DFT2 mutation rates. Mutation rates were estimated using linear regression except for “Substitutions (BEAST)”, which was estimated using a Bayesian phylogenetic approach (*27*). Rate ranges represent 95% confidence interval of the linear fit except for “Substitutions (BEAST)”, where range represents 95% Bayesian credible interval. DFT1 hypermutator clade E was excluded from substitution and indel rate calculations. Ratio ranges represent error-propagated 95% confidence intervals. Level of significance of F-test for linear fit is shown, ratios of mutation classes which did not show significant linear fits are not displayed. *, *F*-test *p* < 0.01. **, *F*-test *p* < 1 × 10^-08^. n.s., not significant.

|  | DFT1 rate, per year | DFT2 rate, per year | DFT2:DFT1 rate ratio |
| --- | --- | --- | --- |
| Substitutions (BEAST) | 215.5 [212.2-218.5] | 516.7 [505.8-527.7] | 2.397 [2.336-2.457] |
| Substitutions (regression) | 163.2 [119.6-206.8]** | 496.3 [436.0-556.5]** | 3.041 [2.753-3.329] |
| SBS1 substitutions | 12.6 [10.5-14.7]** | 17.0 [13.1-20.9]** | 1.351 [1.074-1.629] |
| SBS5 substitutions | 150.6 [108.8-192.4]** | 479.2 [422.1-536.4]** | 3.182 [2.886-3.478] |
| Indels | 22.2 [12.6-31.8]** | 86.5 [73.0-100.0]** | 3.897 [3.445-4.348] |
| ID1 indels | 9.8 [3.5-16.1]* | 13.0 [6.7-19.3]* | 1.326 [0.539-2.113] |
| ID2 indels | 1.4 [-0.43-3.25], n.s. | 27.1 [20.5-33.6]** | - |
| LINE-1 insertions | -0.2 [-0.35-0.04], n.s. | 24.1 [18.5-29.6]** | - |
| Rearrangement events | 1.1 [0.3-1.8]* | 3.6 [2.7-4.5]** | 3.394 [2.646-4.142] |
| Copy number events | 0.7 [0.2-1.1]* | 5.6 [4.7-6.5]** | 8.391 [7.701-9.080] |

The relationship between substitution burden and sampling date is remarkably linear in both DFT1 and DFT2. Nevertheless, a group of DFT1 tumours can be observed with fewer substitutions attributable to both SBS1 and SBS5 than expected (Figure 3G). These tumours belong to a single branch of the phylogenetic tree, clade C2/3, corresponding to the group of clade C tumours sampled in north-west Tasmania (Figure 3G). The mutation rate inferred when considering only these tumours (179 mutations per year, 95% confidence interval 131-227) is similar to that of the remaining DFT1 tumours (202 mutations per year, 95% confidence interval 166-238), however, there are approximately 1,200 fewer mutations genome-wide in the overall clade C2/3 burden than expected. Indeed, clade C2/3 tumours accounted for a significant fraction of the variance in the linear fit for substitutions, attributable to both SBS1 and SBS5, regressed against time (Figure 3G). These observations suggest that a transient reduction in mutation rate occurred during the chain of transmissions taking place between 1991 and 2003 that transported DFT1 into Tasmania’s north-west, perhaps due to a temporary reduction in cell division rate. Such fluctuations in mutation rate may not be uncommon, with detection in this particular case made possible due to the long internal branch and particularly dense sampling of DFT1 clade C2/3.

### A DFT1 hypermutator lineage

Although most DFT1 and DFT2 tumours possess very similar mutational spectra, a single DFT1 tumour, the unique representative of the early divergent clade E, named 377T1, had a highly distinctive pattern of mutations (Figure 3H). Signature fitting suggested that, in addition to SBS1, SBS5, ID1 and ID2, this tumour also carried mutations attributable to mutational signatures SBS6 and ID7 (Figure S9). Furthermore, 377T1 carried six and ten times more substitutions and indels, respectively, than expected from other DFT1 tumours sampled at a similar time (Figure 3H). As SBS6 and ID7, as well as elevated activity of ID1 and ID2, have been linked to deficiencies in DNA mismatch repair (MMR) (*23, 24*), these observations suggest that a clonal ancestor of 377T1 lost MMR function. In order to identify the lesion that disrupted MMR in 377T1, we screened the sequences of genes encoding selected MMR effectors in DFT1 tumour genomes, and discovered a focal deletion specific to 377T1 that removed a single copy of *MLH1* (Figure 3H). Supporting a role for this gene, the 377T1 mutational spectrum is highly reminiscent of that reported in human cells lacking *MLH1* (*28*). No mutations, however, were detected in the remaining copy of *MLH1,* and we speculate that this may have been transcriptionally silenced, for example by promoter DNA methylation.

### Transposable element activity in DFT1 and DFT2

Transposable elements are frequently active in human cancer (*29*), but it is not known if these are mobilised in Tasmanian devil cancers. Several families of transposable elements are annotated in the new reference genome, mSarHar1.11, including 1,948 full-length long interspersed nuclear element 1 (LINE-1) retroelements (Table S1). We systematically screened for somatic LINE-1 insertions in DFT1 and DFT2 and found high LINE-1 transposition activity in DFT2, with hundreds of insertions detected. In DFT1, however, no clear evidence of LINE-1 activity was found (Table S4). LINE-1 mobilisation events were observed throughout the DFT2 phylogenetic tree and accumulated linearly with time (Figure 4A, Table 2, Table S4).

**Fig. 4:**
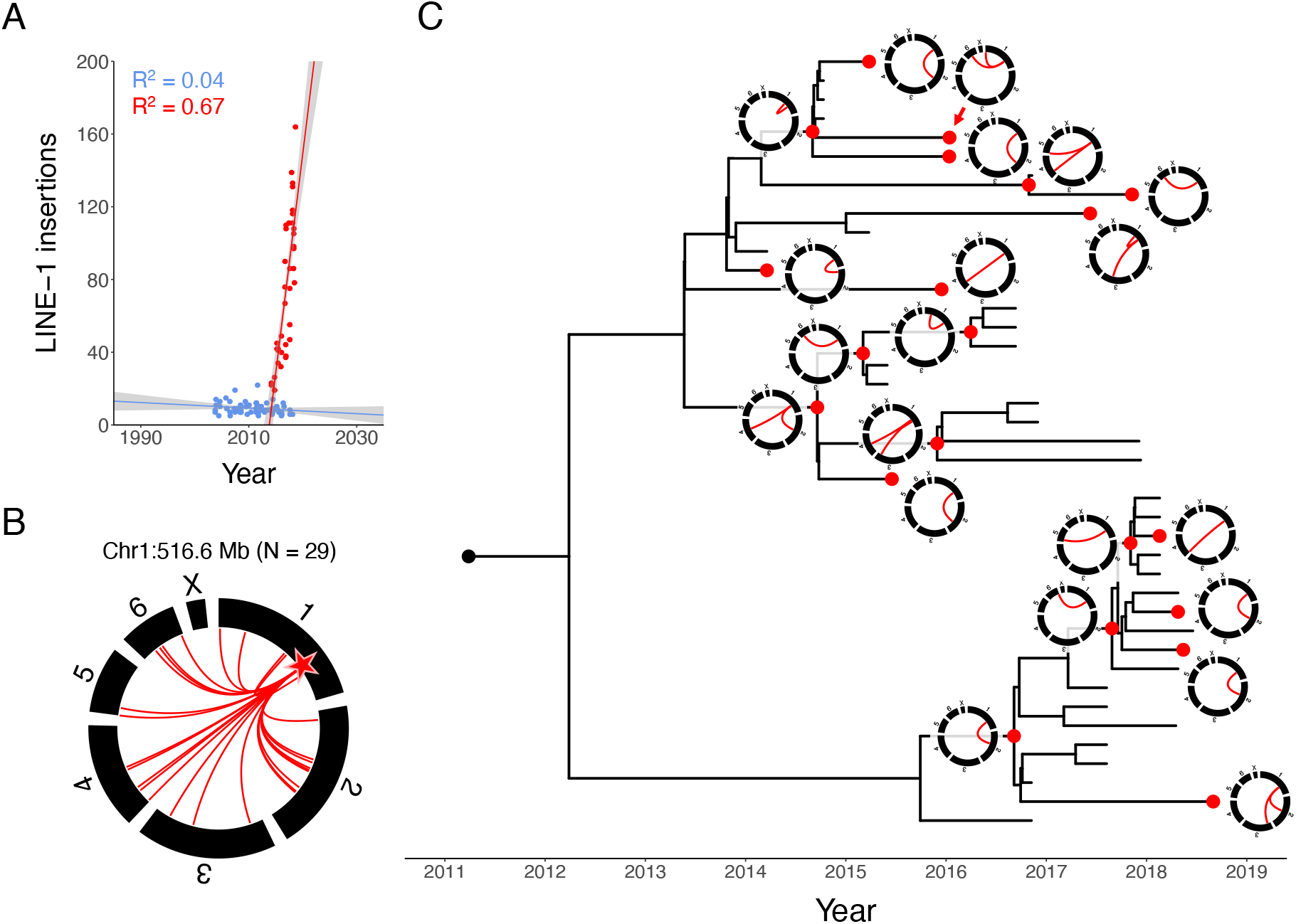
LINE-1 transposable element activity in DFT1 and DFT2. (**A**) Rate of LINE-1 insertion accumulation in DFT1 (blue) and DFT2 (red). Each point represents a tumour, plotted by sampling date. Lines represent linear regression, grey shading 95% confidence interval. (**B**) DFT2 3’ transduction activity of a LINE-1 source element at chromosome 1:516.6 megabases (Mb) (star). In the circos plot chromosomes are represented by black bars, and red arcs connect source element to 3’ transduction integration site. (**C**) DFT2 phylogenetic tree with circos plots illustrating temporal activity of the LINE-1 source element located at chromosome 1:516.6 Mb. Nodes corresponding to each circos plot are represented in red. Source data available in Table S4.

Transcriptional read-through occasionally mobilises genomic DNA downstream of LINE-1 source elements in a process known as 3’ transduction (*29*). A subset of DFT2 LINE-1 insertions carried 3’ transductions, identifying 35 functional LINE-1 source elements in DFT2 (Table S4). Although most DFT2 source elements could be associated with only a single LINE-1 3’ transduction event, one source element located on chromosome 1 spawned at least 29 LINE-1 3’ transductions, with activity continuing throughout the DFT2 phylogenetic tree (Figures 4B and 4C). Overall, these findings reveal that LINE-1 retroelements are transposition-competent in Tasmanian devil genomes, and that their activity varies substantially between DFT1 and DFT2.

### Genome rearrangement in DFT1 and DFT2

The availability of mSarHar1.11 enabled detailed reconstruction of the chromosomal rearrangements that initiated DFT1 and DFT2. The genome catastrophe that marked the origin of DFT1 is focused on the tip of the long arm of chromosome 1 (*10, 14, 30*). This region is massively internally rearranged through dozens of inversions interspersed with short deletions and interchromosomal translocations (Figure 5A, Tables S5 and S6). These changes are compatible with a complex chromothripsis event, as previously proposed (*14*). The early rearrangements of DFT2 are less clustered than those of DFT1 (Figure 5A, Table S5 and S6) (*10*). Chromosome ends are notably involved in rearrangement in both DFT1 and DFT2, consistent with a role for telomere dysfunction in DFT initiation (*10, 14, 30*).

**Fig. 5:**
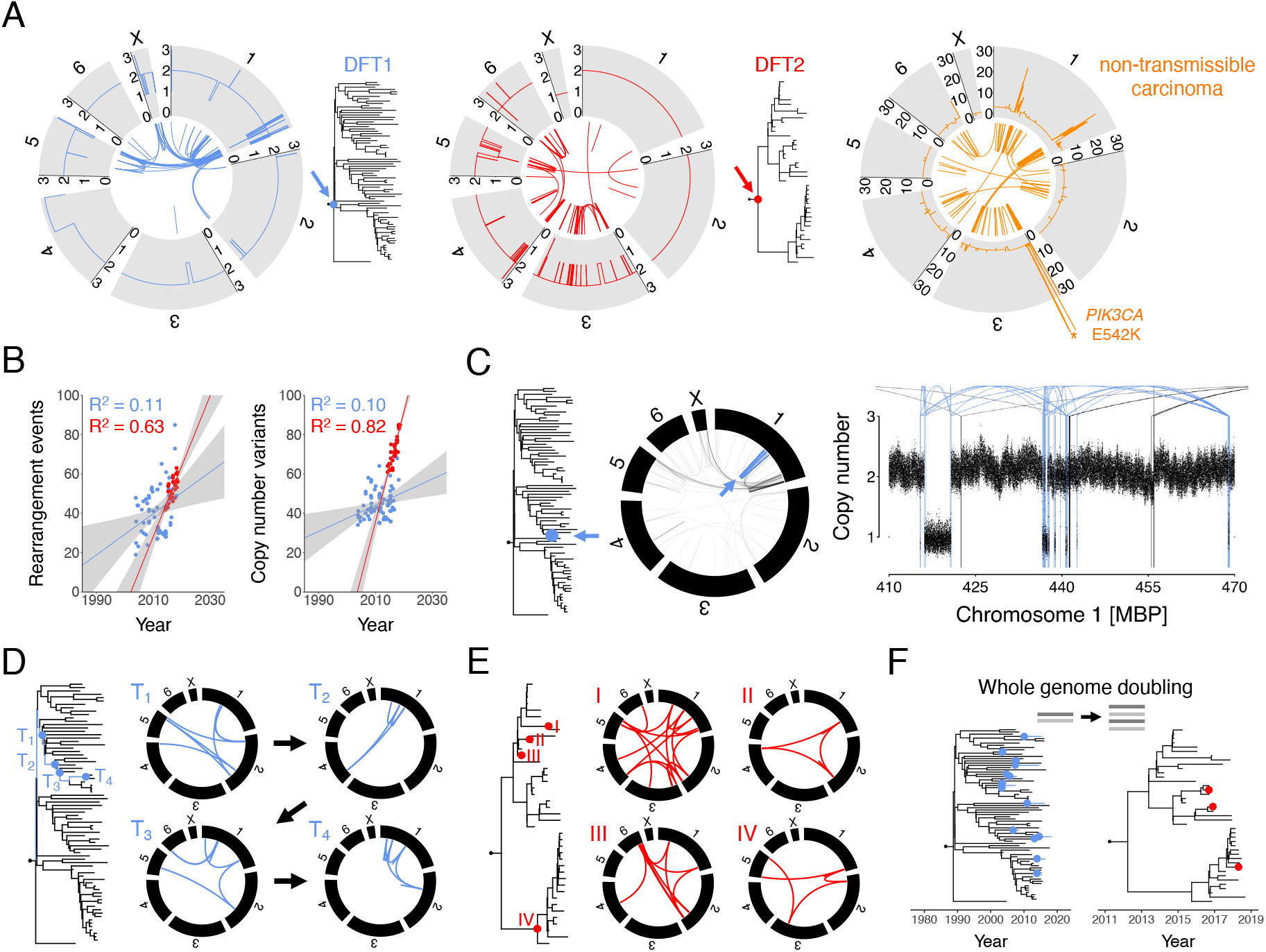
Genome rearrangement in DFT1 and DFT2. (**A**) Rearrangement and copy number profiles of the DFT1 (left, blue) and DFT2 (centre, red) most recent common ancestor tumours (trees, arrows). Chromosomes are represented by grey blocks annotated with copy number state. Inner arcs represent rearrangements. Right, rearrangement and copy number profiles of a single Tasmanian devil non-transmissible carcinoma. The location of the highly amplified E542K mutation in *PIK3CA* is labelled (asterisk). (**B**) Rates of accumulation of rearrangement events (left; “events” denotes that clustered rearrangements have been merged) and copy number variants (right) in DFT1 (blue) and DFT2 (red). Tumours are represented by points, plotted by sampling date. Lines represent linear regression, grey shading 95% confidence interval. (**C**) Example of a late chromothripsis event in DFT1. A single DFT1 tumour (blue dot, arrow on phylogenetic tree) carries a chromothripsis event on chromosome 1; on the circos plot rearrangements unique to the affected tumour are drawn in blue and shared rearrangements that were acquired prior to this tumour’s divergence are drawn in black. Right, copy number plot illustrates rearrangements involving the chromothriptic region (arcs; blue arcs are unique to this tumour, black arcs are shared with other tumours), and copy number is illustrated with binned coverage; each bin represents normalised read coverage in a 1 kb window. MBP, megabase pairs. (**D** to **E**) Examples of chromoplexy events in DFT1 (left) and DFT2 (right). In both cases, positions of nodes represented by each circos plot are illustrated on the relevant phylogenetic tree, either along a four-step time-resolved (T_1_ - T_4_) branch trajectory in DFT1 (**D**) or throughout the DFT2 phylogeny (**E**). Chromosomes are represented by black blocks, rearrangements by coloured arcs. (**F**) Timing of whole genome doubling events in DFT1 (15 events) and DFT2 (3 events). Estimated date of each whole genome duplication is illustrated on tree with coloured dot. Further information and source data are available in Figures S10 and S11, and Tables S5 and S6.

The genome of the spontaneous non-transmissible anal sac carcinoma showed dramatic rearrangement and copy number alteration (Figure 5A, Table S5 and S6). This cancer’s pattern of stepwise amplification is compatible with the activity of several breakage-fusion-bridge cycles. It is notable that the copy number landscape of this tumour is significantly more complex than those of the respective most recent common ancestors of DFT1 and DFT2, indicating that, just as in humans, there are several routes to carcinogenesis in Tasmanian devils.

Rearrangement events and copy number variants (CNVs) both accumulated linearly with time in DFT2 (Figure 5B, Table S5 and S6). Although slight temporal increases were detected in DFT1, these were only marginally significant, confirming previous findings that the rate of genomic structural change in DFT1 is barely detectable above background variation among sublineages (*18*). Despite this, it is noteworthy that the group of DFT1 clade C2/3 tumours that carried fewer substitution mutations than expected (see Figure 3G) also showed fewer rearrangement events and copy number variants (Figure S10), suggesting that the transient reduction in mutation rate occurring on the westward transmission chain operated across mutation classes.

The spectra of polymorphic (i.e. occurring after each lineage’s most recent common ancestor) genomic rearrangements in DFT1 and DFT2 were similar, with small-scale alterations dominating (Table S5 and S6). Several more complex events were also observed in both lineages, however, including occasional chromothripsis (Figure 5C) and ongoing chromoplexy (Figures 5D and 5E). We investigated the genomic contexts and haplotype specificity of a subset of CNVs observed to occur repeatedly either within or between DFT lineages (*18*); one of these was associated with repetitive structural features likely triggering genome instability (Table S6). Copy-neutral variation in minor copy number was rare in DFT1 and undetectable in DFT2, consistent with these tumours’ overall patterns of copy number stability (*18*).

### Whole genome doubling in DFT1 and DFT2

Among the 78 DFT1 and 41 DFT2 tumours analysed, 16 DFT1s and 3 DFT2s were identified as likely tetraploid, defining 15 DFT1 and 3 DFT2 whole genome duplication events. By counting the number of substitution mutations occurring prior and subsequent to genome duplication in each tetraploid lineage, and applying the previously inferred substitution mutation rates, we estimated the dates upon which genome doubling occurred. This identified whole genome duplications that predated sampling of tumours by up to 7 years (median 1.8) (Figure 5F, Figure S11, Table S6). DFT tumours that had undergone genome duplication showed an increased frequency of whole-chromosome or whole-chromosome-arm gain or loss events, compared with diploid tumours (Fisher’s exact test *p* < 0.01, Table S6). This may at least in part be due to mitotic spindle defects introduced secondary to centrosome duplication (*31*), or due to a shortage of chromosome replication effectors in the first cell cycle following genome doubling (*32*); alternatively, it is possible that such large-scale aberrations are better tolerated in the tetraploid state.

### Signals of selection in DFT1 and DFT2

The mutations that initiated DFT1 remain unknown, although a number of candidates have been proposed (*10, 11, 30*). It seems almost certain that the catastrophic event at the origin of DFT1 produced one or more driver mutations. The complex disruption of a single copy of *LZTR1* (*30*) is the most plausible driver candidate associated with this event (Figure 6A and 6B). In DFT2, focal copy number amplification of *PDGFRA* is shared by all DFT2 tumours and remains a strong early driver candidate (Figure 6A) (*10*). In contrast to DFT1 and DFT2, the non-transmissible carcinoma carries recognisable driver mutations in well-characterised cancer genes (E542K *PIK3CA* mutation amplified to more than sixty copies; *TP53* truncation; *NOTCH2* mutations) (Table S6, Table S7; see Figure 5A). Overall, the paucity of clear early driver mutations in DFT1 and DFT2, as well as the absence of causative cancer genes shared by both lineages, suggests that these cancers arose from a cell type that, perhaps by virtue of its epigenetic or transcriptional state, was predisposed to carcinogenesis, requiring only minimal genetic perturbation in order to produce transmissible cancer.

**Fig. 6:**
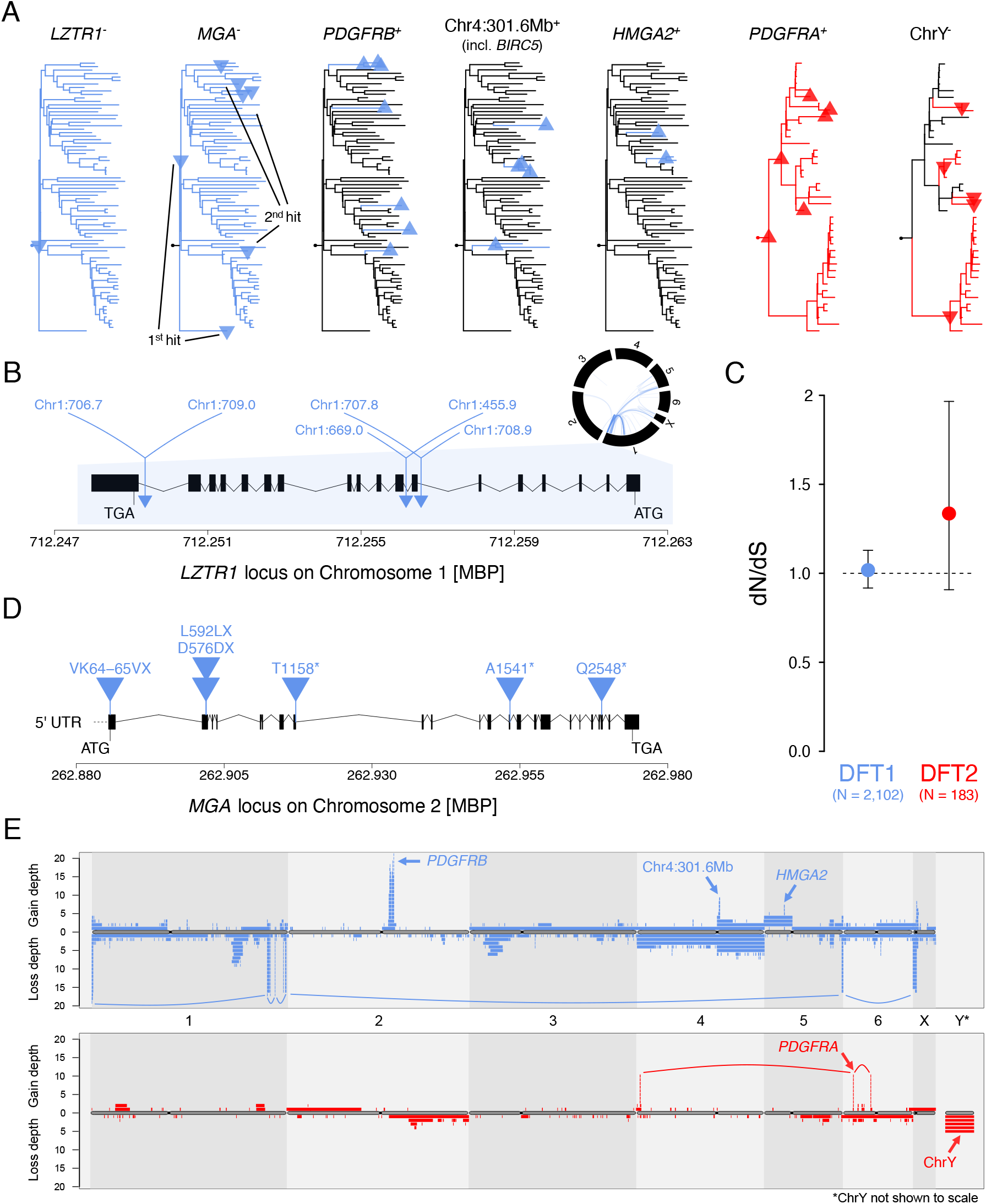
Signals of selection in DFT1 and DFT2. (**A**) Phylogenetic positions of candidate driver mutations in DFT1 (blue) and DFT2 (red). Upward-pointing triangles and “+” notation represent copy number amplifications; downward-pointing triangles and “-” notation represent copy number losses or gene inactivation events. Multiple gains or losses in the same phylogenetic node are only represented once. (**B**) Rearrangement of a single copy of *LZTR1* in DFT1. *LZTR1* (exons represented by black boxes, introns with black connectors) occurs within the densely rearranged region of chromosome 1 that is common to all DFT1s (circos plot; black bars represent chromosomes and blue arcs represent rearrangements common to all DFT1s; Table S5). The location of each rearrangement in *LZTR1* is represented by a triangle, with the coordinates of each partner locus labelled. MBP, megabase pairs. (**C**) Normalised ratio of nonsynonymous-to-synonymous substitutions and indels (dN/dS) in DFT1 and DFT2. Dashed line indicates dN/dS=1 (neutrality) and bars represent 95% confidence intervals. (**D**) Genomic representation of the *MGA* locus on chromosome 2 in DFT1, exons represented by black boxes, introns with black connectors. Blue triangles represent the six coding mutations identified in this gene, all of which are truncating (Tables S7 and S8). MBP, megabase pairs; 5’ UTR, 5’ untranslated region. (**E**) Map representing copy number variants (CNVs) detected within the sampled cohort of 78 DFT1 (upper, blue) and 41 DFT2 (lower, red) tumours. Chromosomes are represented horizontally, with chromosome Y not shown to scale. Each CNV is represented by a coloured bar, with copy number gains illustrated above the grey chromosome representation (“gain depth”) and copy number losses illustrated below the chromosome representation (“loss depth”). Mitotically inherited CNVs are represented once, thus each coloured bar represents a unique CNV occurrence. CNVs that co-occur in the same tumours, and are thus likely to be linked, are connected with coloured arcs; in DFT1, the set of linked losses are associated with the unstable small chromosome known as “marker 5” (*18*). Arrows label candidate driver genes or genomic coordinates associated with prominent focal amplicons. Data associated with figure are available in Tables S6-S8. Table S6 shows haplotype phasing of selected recurrent CNVs.

To explore ongoing evolution in DFT1 and DFT2, we first used *dNdScv* (*33*) to analyse evolutionary signal among substitution and indel mutations (Figure 6C, Table S8). This provided no evidence for widespread negative selection acting to remove deleterious mutations from the coding genomes of DFT1 or DFT2. However, a single gene in DFT1, *MGA*, which encodes a transcription factor that opposes *MYC* activity, showed plausible signs of positive selection through repeated truncation (global likelihood ratio test *q* < 0.005) (Figure 6D). *MGA* has been implicated in cancer, although its driver status is not confirmed (*34, 35*), and occurs in a haploid state in nearly all DFT1s (Figure 6A).

Next, we searched for evidence of late drivers involving copy number variation. We created a chromosome map displaying total CNV burden within the sampled DFT1 and DFT2 population, and examined this for focal amplification (Figure 6E). This screen detected the previously described repeated amplification of *PDGFRB* in DFT1 (Figure 6A and 6E) (*10, 18*) and indicated that further copy number gains of the early *PDGFRA* amplicon in DFT2 have occurred repeatedly in DFT2 clade A (Figure 6A and 6E). This analysis also identified two known recurrent focal amplifications on chromosomes 4 and 5 in DFT1, the latter containing *HMGA2,* and the former carrying 16 genes including *BIRC5* (*18*). In addition, although they are not recurrent, the focal amplification of *RAC1* to four copies in a single DFT1, and focal homozygous deletion of *PTEN* in one DFT2, stand out as potential late driver events (Table S6).

DFT2 arose from a male founder devil and thus carries chromosome Y. The skew towards male hosts present in the DFT2 population (*7*), as well as a previous observation that chromosome Y had been lost from a single female DFT2 host, prompted speculation that loss of chromosome Y (LoY) may be under positive selection in DFT2 by reducing the immunogenicity of this cancer in female hosts (*10*). We investigated this hypothesis by analysing copy number of chromosome Y in our panel of DFT2 tumours. We detected five LoY events throughout the phylogeny of the 41 DFT2 tumours analysed, one of which occurred in the ancestor of DFT2 clade B and is shared among all tumours of this group (Figure 6A and 6E). Somatic LoY is commonly observed in human normal and cancer cells, and the role of selection in driving this alteration in these contexts is poorly understood (*36–38*). Thus, although suggestive, we cannot confirm that DFT2 LoY is under positive selection; indeed, somatic LoY was observed in the analysed non-transmissible devil anal sac carcinoma (Table S6). However, it is noteworthy that a previous study that tracked the karyotype of a chrY^+^ DFT2 cell line through two hundred passages *in vitro* made no mention of LoY in this immunologically neutral setting (*39*). If the presence of chromosome Y is indeed an immunological barrier to the colonisation of female hosts, then no sex imbalance would be expected among hosts of chrY^-^ DFT2.

## Discussion

The assembly of a highly complete and contiguous reference genome for the Tasmanian devil has enabled comprehensive genomic characterisation of this species’ two transmissible cancers. DFT1 and DFT2 are independent realisations of a common biological phenomenon. Although the two cancers are overall highly similar in their genome features, especially when compared to a non-transmissible Tasmanian devil cancer, several differences exist: this ecological niche will tolerate different forms.

A particularly striking difference between DFT1 and DFT2 is the elevated mutation rate, observable across mutation classes, of DFT2 (Table 2). One explanation for this would be that DFT2 has a faster cell division rate than DFT1, and thus greater opportunity for the accrual of mutations associated with DNA replication. If true, this might influence relative growth rates and generation times of DFT1 and DFT2, with potentially complex epidemiological implications. However, other differences in cell state unrelated to division rate may underlie this observation. Furthermore, although it is tempting to attribute the elevation in rates across different mutation classes in DFT2 to a common cause, it is possible that these are, in fact, unrelated, particularly as the magnitude of difference varies among mutation classes and signatures (Table 2). More generally, the mutation rates inferred from DFT1 and DFT2 provide evidence that large-scale mutations, including rearrangement events, transposon insertions and copy number variants, can have clock-like properties within individual cancers.

Once arisen, mutations become subject to selection. Positive selection, acting to increase frequency of mutations conferring advantageous traits, is usually the dominant force in cancer evolution; negative selection, operating to remove deleterious mutations, is also detectable in cancer, although weak (*33*). In transmissible cancers, the stochasticity of transmission may decrease the efficiency of selection, and neutral processes, such as genetic drift, are likely to be of particular importance in their evolution (*40*). Nevertheless, and despite the small sample size of our study, plausible signals of positive selection were detectable in DFT1 and DFT2, and it is likely that these are operating to increase fitness of cells within tumours (e.g. *PDGFRB* and *PDGFRA* amplification in DFT1 and DFT2, respectively, and *MGA* loss-of-function in DFT1) and to enhance transmission potential (e.g. chromosome Y loss in DFT2). Genetic variants that increase somatic mutation rate are themselves often causatively involved in cancer through their tendency to predispose cells to acquisition of secondary adaptive mutations. This may be exemplified in the putatively positively selected heterozygous truncating mutation in *MGA* observed in MMR-deficient DFT1 clade E.

Predicting the future dynamics and impacts of DFT1 and DFT2 requires knowledge of these diseases’ epidemiological parameters. Although estimates of basic reproductive number (*R*_0_) and generation time have been proposed for DFT1 (*15*), considerable uncertainty remains. Phylodynamics methods provide tools for inference of epidemiological metrics from pathogen genomes; however, the small sample size and geographical structuring of our tumour data set make it unsuitable for such analysis (*41,* but see *42*). While we cannot predict the evolutionary outcomes of DFT1 and DFT2, one observation that is worthy of comment is the surprisingly long delay between the origin of DFT1 (1982-1989) and its detection (1996). During this interval several hundred devils were examined in north-eastern Tasmania, the location of DFT1’s first observation, but no evidence of DFT was recorded (*4*). This suggests that DFT1 may have remained at low frequency during this time, and is compatible with a relatively low *R*_0_, or a longer than expected generation time. This observation, together with that of the superspreading event that occurred shortly after DFT1’s origin which involved transmission of a tumour from a single donor to at least six recipients and founded the six DFT1 clades, lends credibility to the hypothesis that *R* may be over-dispersed in DFT, and that a large fraction of transmissions may funnel through a small number of infectious tumour donors (*43*). Tumour, host and seasonal factors may influence individual transmission potential (*44*).

DFT1 and DFT2 have revealed the existence of a biological niche suited for transmissible cancers in Tasmanian devils. There is no evidence that these cancers emerged as a direct consequence of human actions through, for example, the introduction of chemical carcinogens or oncogenic viruses. Thus, it seems most likely that DFTs are a natural part of Tasmanian devil ecology. Although postcolonial human activities may have created conditions that indirectly benefitted DFT emergence or spread, for example through habitat modification that may have supported increased devil density (*45*), it is very likely that DFTs have occurred in the past, and that additional clones will emerge in the future. Notably, many incipient DFTs may die out before detection, particularly if these diseases possess superspreading dynamics. While no specific actions can be taken to prevent the establishment of new DFTs, it will be important to continue close monitoring of wild and captive devil populations.

Although DFT transmissible cancers might themselves be natural occurrences, these diseases’ devastating impact on their host species is exacerbated by anthropogenic threats including loss of habitat and roadkill (*46, 47*). Several recent studies have used longitudinal monitoring data to parameterise models predicting future Tasmanian devil population size, and have argued against DFT1-induced extinction as a likely outcome (*48–50*). However, there is consensus that the species remains under threat, particularly given that its potential for persistence at much reduced density is unknown. It is thus important that adaptive monitoring, research and management continue to be prioritised to ensure long term conservation and resilience of the Tasmanian devil (*46*).

Overall, this survey of the genomes of the two Tasmanian devil transmissible cancers has illuminated the evolutionary history of these unusual pathogens. Our analysis suggests that Tasmanian devils host a cell type that is poised for transmissible cancer transformation, with only minimal somatic genetic disruption required for these to be unleashed. Once established, DFT clones continue to acquire mutations at remarkably constant rates and, although the majority of these are neutral, a small subset drive further adaptation to the niche. The future trajectories of DFT lineages and their Tasmanian devil hosts remain uncertain; however, this study provides a vantage point from which to further explore the evolution and impacts of transmissible cancers in this iconic marsupial species.

## Acknowledgements

We are grateful to Ben Lehner, Hannah Siddle, Rachel Owen, Annalisa Gastaldello, Andrew Flies, Kate Hughes, Simon Mayes, Francesca Giordano, Ian Mickleburgh, Aylwyn Scally, Bronwen Aken, the Wellcome Sanger Institute sequencing pipelines team, and past and current members of the Transmissible Cancer Group for technical assistance and helpful discussions. We also thank students, staff and volunteers involved in Tasmanian devil field work and sample collection. This work was supported by a grant from Wellcome (102942/Z/13), as well as Eric Guiler Tasmanian Devil Research Grants from the University of Tasmania Foundation. Additional field work was supported by grants from the Australian Research Council (LP0561120, LP0989613, DP110102656 – MEJ; DE170101116, LP170101105 – RH), the US National Science Foundation (DEB-1316549 – MEJ and EPM), and the US National Institutes of Health (1R01GM126563-01 – MEJ). MRS was supported by a Gates Cambridge Trust Scholarship. For the purpose of open access, the authors have applied a Creative Commons Attribution (CC BY) license to any Author Accepted Manuscript version arising.

## Supplementary Materials for

### 1. Tissue sampling and animal ethics

Tissues were sampled from: wild Tasmanian devils that were captured and subsequently released; animals found dead; or animals euthanized for welfare reasons. All animal procedures were performed under a Standard Operating Procedure approved by the General Manager, Tasmanian State Government Department of Natural Resources and Environment (NRE), in agreement with the NRE Animal Ethics Committee, or were approved under University of Tasmania Animal Ethics Permits A009215, A0010296, A0011436, A0011696, A0012513, A0013326, A0013685, A0014976, A0015835, and A0016789, with State Scientific Permits TFA 08211, TFA 12200, TFA 14228, TFA 15214, TFA 17176, TFA 18028 and TFA 19144. Sample collection procedures were approved by the University of Cambridge Department of Veterinary Medicine Ethics and Welfare Committee (CR191). Recorded trapping locations were displayed using *ggmap* v3.0.0 (*52*).

### 2. mSarHar1.11 genome assembly and annotation

A new Tasmanian devil reference genome, mSarHar1.11, was assembled using DNA sequences from 91H, the Tasmanian devil fibroblast cell line used in the previous Tasmanian devil genome assembly, DEVIL7.0 (*13*). The female donor individual was captive bred (*13*).

#### 2.1 Data generation for mSarHar1.11 genome assembly

##### Oxford Nanopore Technology sequencing

High molecular weight DNA was extracted using the Qiagen Genomic Tip kit (Qiagen, Hilden, Germany) and used to prepare Oxford Nanopore Technology ligation (LSK108, LSK109) and rapid (RAD004) sequencing libraries. In addition, ultra-high molecular weight DNA was extracted using a low-shearing DNA extraction protocol and used to generate RAD004 rapid sequencing libraries (*19*). Libraries were sequenced on GridION and PromethION instruments (standard libraries: 76.40x whole genome coverage, read N50 9.1 kilobases (kb); ultra-long sequence libraries, 11.86x whole genome coverage, read N50 57.1 kb). Nanopore sequencing details are summarised in Table S1.

##### 10x Chromium linked-read sequencing

Chromium linked-read libraries with insert size 300-500 base pairs (bp) were prepared from high molecular weight DNA according to manufacturer’s instructions (10x Genomics, Pleasanton, USA). These were sequenced with 150 bp paired-end reads on two Illumina HiSeq XTen lanes (Illumina, San Diego, USA) to ∼60x depth, with a linked-read N50 of 184.1 kb.

##### Hi-C sequencing

Hi-C libraries were prepared using the Dovetail Hi-C kit according to manufacturer’s instructions (Dovetail Genomics, Scotts Valley, USA). Libraries with 350 bp insert size were sequenced with 150 bp paired-end reads using two lanes of an Illumina HiSeq XTen instrument (Illumina, San Diego, USA), yielding approximately 190 Gb (∼60x genome coverage).

##### Bionano optical mapping

High molecular weight DNA was isolated using the IrysPrep Plug Lysis Long DNA Isolation Protocol, and molecules labelled according to manufacturer’s instructions (Bionano Genomics, San Diego, USA). Molecules were imaged using a Bionano Irys instrument (Bionano Genomics, San Diego, USA). Several cycles were performed to reach an average raw genome depth of ∼30x.

#### 2.2 mSarHar1.11 genome assembly

##### Initial contig assembly

Initial contig assembly with Oxford Nanopore Technology reads was performed using wtdbg (v1, https://github.com/ruanjue/wtdbg) with options “-H -k 21 -S 1.02 -e 3 2”. This produced a 3.11 Gb raw assembly with 530 contigs (N50, 63.9 megabases (Mb); N90 9.26 Mb). Assembly algorithm benchmarking is summarised in Table S1.

##### Contig polishing

10x Chromium linked-reads were aligned to assembled contigs using Long Ranger (https://github.com/10XGenomics/longranger; https://support.10xgenomics.com/genome-exome/software/downloads/latest). Two runs of polishing were carried out using Freebayes (*53*), and a polished consensus contig file obtained.

##### Chromosomal scaffolding

Contig scaffolding was performed with 10x Genomics linked reads using scaff10x (https://github.com/wtsi-hpag/Scaff10X) and subsequently with Hi-C reads using scaffHiC (https://github.com/wtsi-hpag/scaffHiC). Unclosed gaps were filled with poly-N stretches of length 200 bp in the Hi-C guided assembly.

##### Final curation of mSarHar1.11

We followed genome assembly curation procedures as previously outlined (*54*) using gEVAL (*55*). Bionano optical mapping data were aligned along with 10x linked reads and Hi-C reads. Assembly errors, haplotypic duplicated contigs and contaminating sequences were removed. Several contigs were merged, reducing the number from 530 to 445. This process yielded an assembly composed of seven major chromosomal scaffolds containing 3.09 Gb assembled sequence (scaffold N50, 611 Mb) and 97 unassigned contigs containing 5.1 Mb (Table S1).

##### Mitochondrial genome and chromosome Y

The DEVIL7.0 mitochondrial genome, which was derived from the same 91H donor individual (*13*), was lifted over to mSarHar1.11 as a separate contig. We also added two contigs derived from chromosome Y of a male individual which had been previously sequenced and assembled (202H1 (*10*)). These contigs carry chromosome Y-encoded genes *SRY* and *KDM5D,* and contribute a total length of ∼130 kb.

##### Chromosome 1 and chromosome 2 nomenclature

There are two nomenclature systems in use for Tasmanian devil chromosomes, which differ with respect to the naming of the two largest chromosomes (*1, 56*). The chromosome labelled “1” in the first system is named “2” in the second, and vice versa. In mSarHar1.11, labels of chromosome 1 and chromosome 2 reflect those originally assigned by Martin and Hayman (*56*). This differs from DEVIL7.0, which used the system adopted by Pearse and Swift (*1*).

##### Centromere locations and chromosome orientation

Centromere positions were inferred from the mSarHar1.11 sequence’s Hi-C contact map, and independently assessed against a set of devil chromosome arm-specific marker genes (*14*). Centromere coordinates are summarised in Table S1.

#### 2.3 mSarHar1.11 genome annotation

Genome annotation was performed using the Ensembl genome annotation pipeline (*20*) with steps outlined below, as well as the automated NCBI RefSeq annotation pipeline (*57*).

##### Repeat annotation and masking

Prior to annotation, the genome was repeat-masked with RepeatMasker v4.0.5 (https://repeatmasker.org/) using the crossmatch engine and the Repbase vertebrates v2017-0127 library (*58*). This masked 49.6 percent of the genome. In addition, we also built a custom library generated with RepeatModeler v1.0.11 (*59*), and combined this with RepeatMasker to generate a secondary repeat track, masking 52.97 percent of the genome. Additional repeats were generated using Dust v1.0.0 (*60*) with default settings for low complexity regions and Tandem Repeat Finder (*61*) with default settings for tandem repeats. For downstream gene annotation, the combination of the RepeatMasker repeats derived from the vertebrate library along with the low complexity regions identified by Dust were used to mask the genome for software requiring a softmasked genome as input. Annotations of the repeat features generated from the different approaches were made available through the Ensembl genome browser through a variety of repeat tracks.

##### Transcriptomic annotation

RNA was extracted from Tasmanian devil tissues stored in RNAlater (Sigma-Aldrich, St. Louis, USA) using the AllPrep DNA/RNA/miRNA Universal kit (Qiagen, Hilden, Germany) (Table S1, ENA project accession PRJEB34650). Ribosomal RNA was depleted using Ribo-Zero (Illumina, San Diego, USA), cDNA was synthesised using random hexamer priming, and libraries with insert size 100-300 bp were generated. Three libraries were pooled per lane of a HiSeq 4000 instrument with V4 chemistry (Illumina, San Diego, USA), producing 90-145 million (median 112 million) 75 bp paired-end reads per sample.

These data, along with additional publicly available transcriptome data (*12, 62, 63*), were downloaded from the European Nucleotide Archive (https://www.ebi.ac.uk/ena/) (ENA project accessions: PRJEB28680, PRJNA381841 and PRJNA274196). Reads were aligned to mSarHar1.11 using BWA v0.7.17-r1188 (*64*), with the ‘aln’ command, with a tolerance of 50 percent mismatch to allow for intron identification via split read alignment, and all other settings at default. Reads that were likely to span an intron based on the initial BWA alignment were then realigned in a splice-aware manner using Exonerate v2.2.0 (*65*) with the maximum intron size set to 200 kb and all other settings at default. An Ensembl in-house transcript reconstruction code was then used to build transcript models. A second approach, using STAR v2.5.3a (*66*) to align the reads (with 2-pass mode enabled) and Scallop v0.10.5 (*67*) with ‘-- min_flank_length = 10’ to reconstruct transcript models, was also employed.

Putative transcript structures were examined for coding potential. The longest open reading frame was calculated for each, and the resulting translation was searched against UniProt protein existence level 1 and 2 vertebrate proteins (release 2019_11 (*68*)). Coverage against the database protein and percent identity was calculated. Transcriptomic models created this way formed the primary basis for the annotation of protein-coding and lncRNA genes in the final gene set.

In addition, RNA sequencing reads derived from DFT1 and DFT2 samples from ENA projects PRJEB28680 and PRJNA416378 were downloaded and processed with the same pipeline to create secondary tracks in the browser. These tumour-specific transcript models were not used as part of the main gene set.

##### Homology annotation

We generated a pairwise whole genome alignment against the human GRCh38 assembly using LastZ v1.04.0 (https://www.bx.psu.edu/~rsharris/lastz/), using ‘T=1 K=5000 L=5000 H=3000 M=10 O=400 E=30 --ambiguous=iupac’ with other settings at default. Aligned genomic regions were used to map protein-coding exons from the human GENCODE gene set ((*69*), mapped from Ensembl release 98). For each protein-coding gene in human, coordinates of protein-coding exons were projected through the alignment, with adjustments for potential frameshifts. In addition, we aligned all mammalian proteins from UniProt with protein existence level 1 or 2 to mSarHar1.11 in a splice-aware manner using GenBlast v1.0.139 (*70*), with the settings ‘-g T -c 0.5 -r 10 -i 30 -rl 5000’. For each transcript set (that projected from human, and that generated by aligning UniProt proteins) we calculated the coverage of the original sequence by the new translation and the percent identity between the aligned sequences to one another.

Homology-based models were used to gap fill gene annotations where the transcriptomic data were absent for a particular locus (for example, genes expressed in a particular tissue or development stage not covered by the available transcriptomic data). Additionally, homology-based models were used to supplement or replace models derived from the transcriptomic data in cases where the transcriptomic data appeared to represent a fragmented or short-form transcript.

##### Immunoglobulin and T cell receptor genes

Immunoglobulin (IG) and T cell receptor (TCR) genes were manually annotated. Four steps were taken to identify V, D, J, and C segments of immunoglobulin (IG) and T cell receptor gene loci. (1) BLAST (*71*) searches were performed to identify C, V and J segments using koala, opossum, and human TCR and IG coding sequencings. (2) The recombination signal sequence (RSS) with 23 bp or 12 bp spacers flanking V, D, and J segments were extracted from the koala genome to generate a set of profile hidden Markov models (HMMs) for each TCR or IG locus using HMMER v3.3 (*72, 73*). HMM searches were then performed to identify potential RSSs associated with V, D, and J segments in the new devil genome mSarHar1.11. (3) Species-specific profile HMMs of the RSS were built for devil TCR and IG gene segments by extracting RSSs found in steps 1 and 2. This was followed by a second round of HMM searches with higher sensitivity than those done in step 2. Newly found RSSs were used to update the profile HMMs, after which a final round of HMM searches were performed. (4) Based on all the RSSs found, associated V, D, and J segments were identified upstream, between, and downstream of the relevant type of RSS, respectively.

##### lncRNA annotation

Multi-exon transcript structures built from the short-read data that were over 200 bp sequence length and had no BLAST (*71*) hit, using blastp v2.2.30+ at default settings against the UniProt vertebrate PE1/2 database, to the longest ORF were considered as candidate lncRNAs. Any candidates with an overlapping protein alignment, either from the cross-species protein alignments or projection of coding exons from human, were removed.

##### Pseudogene annotation

Genes with evidence of frame-shifting, or those embedded within repeat-rich genomic regions, were annotated as pseudogenes. Single-exon retrotransposed pseudogenes were identified by aligning candidate transcript sequences against all multi-exon transcripts from the other genes in the gene set using blastn at default settings (*71*). Cases where the single-exon transcript covered 80 percent or more of a multi-exon transcript elsewhere in the genome were classed as retrotransposed pseudogenes.

##### Small non-coding genes

Small non-coding genes were annotated using Rfam v14.0 (*74*) and miRBase v22.1 (*75*). Rfam covariance models were searched against the genome using cmsearch v1.1.2 (*76*), using ‘– ga’ and all other settings at default, and passed through a series of filters to remove likely false positives. miRBase entries were searched against the genome using BLAST (Altschul et al., 1990), before being passed through a series of filters including alignment quality, repeat coverage and minimum fold energy. Structures for the miRNAs were then calculated using RNAfold v2.3.3 (*77*).

##### Data availability

The Ensembl gene annotation for devil was released in Ensembl version 102, including orthologs, gene trees, and whole-genome alignments against other species. We also provide tissue-specific tracks for the RNAseq transcript models, the alignment coverage and complete set of splice junctions identified by our pipeline. Data for the Tasmanian devil genome annotation can be accessed in Ensembl in a variety of ways including the genome browser, the Perl and REST APIs, the ftp site, BioMart and direct access to MySQL databases.

### 3. Sample processing and sequencing

#### 3.1 Whole genome sequencing

Genomic DNA was extracted from devil tumour and normal tissues stored in RNAlater (Sigma-Aldrich, St. Louis, USA) using either the DNeasy Blood and Tissue Kit (Qiagen, Hilden, Germany), AllPrep DNA/RNA/miRNA Universal Kit (Qiagen, Hilden, Germany) or Genomic-Tip Kit (Qiagen, Hilden, Germany). Standard whole genome sequencing libraries with a median insert size of 467 bp were generated and sequenced with 150 bp paired-end reads on an Illumina XTen instrument (Illumina, San Diego, USA). Tumours were sequenced to 83.15x median coverage and normal samples to 40.89x median coverage. Sequence reads from newly sequenced data, as well as those from publicly available data, were aligned to mSarHar1.11 using BWA-mem v0.7.17-r1188 with options -Y and -p specified (*78*). Sample metadata are presented in Table S2.

#### 3.2 Tumour purity calculation

Tumour purity was estimated using two independent methods. The first relied on the offset in read coverage occurring between genomic regions of different copy number states. A 24.7 Mb pericentromeric single copy deletion on chromosome 3 (chromosome 3:192,161,000-216,866,000) is found in all DFT1 and DFT2 tumours examined to date (*10, 18*). Read depth across chromosomes 1-6 was counted in 1 kb bins using bedtools coverage v2.23.0 (*79*). For each tumour, we calculated median coverage across all bins (*C_All_*), as well as median coverage in bins falling within the footprint of the chromosome 3 deletion (*C_Del_*). Purity (ρ) was then calculated as follows:

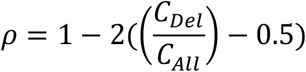

The second approach made use of somatic substitution variant allele fraction (VAF) distributions. VAF distributions were inspected for each tumour, and the value of *VAF*_Het_, corresponding to the maximum density of the heterozygous peak (variants occurring in one of two copies in diploid tumours; or variants occurring in two of four copies in tetraploid tumours) was identified. Purity (ρ) was defined as ρ = 2\**VAF*_Het_. Purity values estimated with both approaches were highly similar (R^2^ = 0.96; Table S2), and estimates using the first method were used for downstream analyses. The non-transmissible carcinoma only had a purity estimate from the second method, however, and this was used in analyses of this tumour.

#### 4. Substitutions and indels

#### 4.1 Substitution and indel calling and genotyping

Substitutions and indels were called and genotyped using Platypus (*80*) with settings as previously described (*10*). Lengths of the new mSarHar1.11 reference chromosomes 1-3 exceed 536 Mb, which requires CSI indexing of BAM and VCF files for downstream processing. While the original Platypus implementation is not fully compatible with CSI indexing, we separately genotyped variants in regions >536 Mb through temporary positional transformations in the input files. Substitutions and indels were filtered as follows (Table S3, Supplementary Data S1):

- *Homozygous-in-reference filter.* Substitutions and indels called with variant allele fraction (VAF) ≥0.9 in the sample corresponding to the reference individual, 91H, were discarded.
- *Strand bias filter.* We rejected variant calls with less than 20% support on either the forward or reverse sequencing strand across all tumour and normal samples.
- *Simple repeats regions filter.* Substitutions and indels lying within a 5 bp window around simple repeat regions, as annotated by Tandem Repeat Finder (*61*), were discarded.
- *Contig and scaffold end regions filter.* We rejected any variant mapping within 500 bp from the start or end of a contig, or within 1000 bp from the start or end of a scaffold.
- *Low and high coverage filter (nuclear genome only).* We excluded any variant with a median coverage of >200 or <10 across 80 normal genomes.
- *Excessive heterozygosity filter.* We removed germline substitutions and indels which, based on their VAF distribution, appeared as heterozygous in ≥85% normal genomes. Table S3 lists sample-specific VAF thresholds used to define heterozygosity.
- *Low-VAF filter.* We removed substitutions and indels which did not meet a sample-specific lower VAF threshold in any tumour. Lower VAF thresholds, listed in Table S3, were defined individually for each tumour after visualising the somatic VAF distribution.

#### 4.2 Variant categorisation

A variant was defined as “present” in a tumour if (1) it was supported by ≥3 reads and exceeded the sample-specific lower VAF threshold outlined above; or (2) it was supported by ≥3 reads and exceeded the sample-specific lower VAF thresholds of ≥50% tumours within the same lineage (i.e. DFT1 or DFT2) carrying the variant with ≥1 supporting read. Variants in 340T, the non-transmissible carcinoma, were defined only using condition (1) above.

A variant was defined as “present” in a normal genome if it was supported by ≥3 reads and exceeded a sample-specific lower VAF threshold. Additionally, variants classed “present” in normal genomes were further defined as heterozygous or homozygous-ALT using sample-specific VAF thresholds (Table S3).

Variants were defined as somatic if they were called as present in ≥1 tumour, but were not identified as present in any normal genome. All remaining variants were classed as germline. The full set of somatic and germline variants, as well as their genotypes in the samples included within this study, are made available (Data S1).

#### 4.3 Ancestral phylogeny

The phylogenetic tree displaying the relationship between DFT1, DFT2 and normal Tasmanian devils, shown in Figure 1C and Figure S2A, was constructed as follows.

Starting with autosomal germline and somatic substitution variants, we identified genomic intervals containing germline copy number polymorphisms (see section 7.1) and removed germline substitutions occurring either within these intervals, or within 10 kb on either side of these intervals. We generated a multiple sequence alignment including those variants called as “present” (see above) in any one of 130 samples (79 normal Tasmanian devils, excluding the reference individual 91H; 38 DFT1 and 12 DFT2 tumours with purity ≥75%; the non-transmissible carcinoma, 340T). In the alignment, the alternative allele was substituted if it was present in either a heterozygous or homozygous state. This alignment included 1,175,235 substitution variants (1,070,436 germline, 104,799 somatic).

A phylogenetic tree was inferred from the alignment using RAxML-NG v.1.0.3 (*81*). One hundred maximum likelihood starting trees steps were used in combination with model parameters GTR+G+ASC_LEWIS. The tree with highest likelihood was displayed using *ggtree* v3.2.0 (*82*).

#### 4.4 Mutational signatures

Somatic substitution mutational spectra were generated as described (*10*), with mutation counts normalised to triplet frequency in mSarHar1.11 (https://github.com/MaximilianStammnitz/SubstitutionSafari), excluding regions excluded from variant calling (see section 4.1). Somatic indel mutational spectra were generated using *Indelwald* (https://github.com/MaximilianStammnitz/Indelwald) without normalisation for genome content.

Signature fitting was performed using single base substitutions signatures from the COSMIC v3.2 database (March, 2021 (*23*)) using *sigfit* (*83*) with four independent 100,000 Markov chain Monte Carlo (MCMC) iterations and 50,000 burn-in cycles, respectively (Table S3).

#### 4.5 Substitution and indel rate analyses

Linear least-squares regression was performed on tumour mutation counts against sampling date using *R* (*84*), excluding tumours without exact sampling dates (458T1, 1439T7, 1524T1, 1525T1). 95% confidence intervals and R^2^ values were calculated using the ‘lm’ function in *R* (*84*). In tetraploid tumours (see section 7.4, Tables S2 and S6), mutation counts were corrected as follows. First, custom variant allele fraction (VAF) thresholds were used to computationally isolate mutations occurring prior and subsequent to whole genome duplication (Table S6). Those occurring after whole genome duplication were corrected for genome opportunity (Table S6).

Signature-specific mutation rate linear regression was done using the same approach, after assigning mutations to signatures as follows. For substitutions, 5’-N[C>T]G-3’ (N, any base) mutations were assigned to signature 1, and remaining mutations assigned to signature 5. For indels, single T insertions at poly-T (≥5 bp) stretches were assigned to signature ID1, and single T deletions at poly-T (≥5 bp) stretches were assigned to signature ID2. In tetraploid tumours, post-tetraploidisation mutations (i.e. those present on 1 of 4, or 1 of 6 copies) were down-sampled, and the remaining mutations assigned to signatures.

#### 4.6 Time-resolved phylogenetic trees

We generated time-resolved phylogenetic trees for DFT1 and DFT2 using tip date sampling implemented in BEAST v1.10.4 (*27*).

##### Somatic substitution pre-processing for timed tree inference

Somatic substitution mutation sets were pre-processed as follows:

1. We randomly subsampled 50% (1 of 4 copies) or 33% (1 of 6 copies) substitutions that occurred after tumour tetraploidisation to account for mutation opportunity (see sections 4.5 and 7.4) (Tables S2 and S6).
2. We subsampled mutations unique to the DFT1 hypermutated tumour, 377T1. The intention was to simulate this sample’s mutation burden under conditions of zero exposure to mutational signature 6. First, we calculated the number of mutations expected in this tumour under such conditions (5,029), given its sampling date, using the linear regression shown in Figure 3C. We then subsampled the mutations called in 377T1 to create a new sample, 377T1-sim. 377T1-sim carried the 1,311 mutations shared between 377T1 and other DFT1s, as well 3,309 mutations unique to 377T1. The latter were sampled from among the 29,462 377T1-unique mutations using a multinomial probability distribution generated using the mutational spectrum presented in Figure 3A.
3. Truncal variants in DFT1 and DFT2, i.e. those occurring before the most recent common ancestor (MRCA) of tumours within each lineage and thus shared by all tumours of the same lineage but absent from the normal devil panel, cannot be definitively assigned as germline or somatic (see further discussion below). After preliminary inspection of maximum likelihood trees, we removed these from the multiple sequence alignments used to generate timed trees, and specified two monophyletic taxa in DFT1 (the first encompassing DFT1 clades A1, A2, B, C and D; the second encompassing DFT1 clade E) and two monophyletic taxa in DFT2 (the first encompassing DFT2 clade A, the second encompassing DFT2 clade B).

Final multiple sequence alignments were constructed for the sets of remaining post-MRCA somatic substitutions specific to DFT1 (N = 171,283) and DFT2 (21,252).

##### Parametrisation of phylogenetic tree inference

We specified a generalised time reversible (GTR) nucleotide substitution model within BEAUTI v1.10.4 (*27*), with base frequency estimation under a gamma site heterogeneity model of four categories. Tree priors were kept at the default ‘coalescent: constant size’, and we set a strict molecular clock. Each tumour’s sampling latitude and longitude (Table S2) were inputted as continuous bivariate trait variables and modelled as Brownian random walks with random jitter of 0.001 (DFT1) or 0.0001 (DFT2) added to overlapping tip coordinates.

For two DFT1 tumours, 458T1 and 1439T7, only the sampling year (2012) could be obtained; in these cases we specified uniform tip date distributions across all 365 days of the calendar year. For two DFT2 tumours, 1524T1 and 1525T1, sampling dates were not available. In both cases we obtained expected sampling dates by extrapolating from observed mutation burdens using linear regression (Figure 3C) (1524T1, 07.11.2017; 1525T1, 08.12.2017). Uniform distributions of 365 days centred on these dates were specified for these tumours.

To counteract ascertainment bias, we inputted the number of invariant sites in the DFT1 and DFT2 alignments, calculated by counting the number of invariant bases (N_A_, N_C_, N_G_, N_T_) in the haploid mSarhar1.11 genome, excluding regions excluded from variant calling (see section 4.1). These corresponded to the following values: DFT1 (N_A_ = 951,704,666; N_C_ = 540,027,583; N_G_ = 539,784,435; N_T_ = 952,062,228); DFT2 (N_A_ = 951,735,820; N_C_ = 540,071,579; N_G_ = 539,827,823; N_T_ = 952,093,054). The input XML file was correspondingly modified as follows (https://groups.google.com/g/beast-users/c/V5vRghILMfw/m/jMtC_DwS5EYJ):

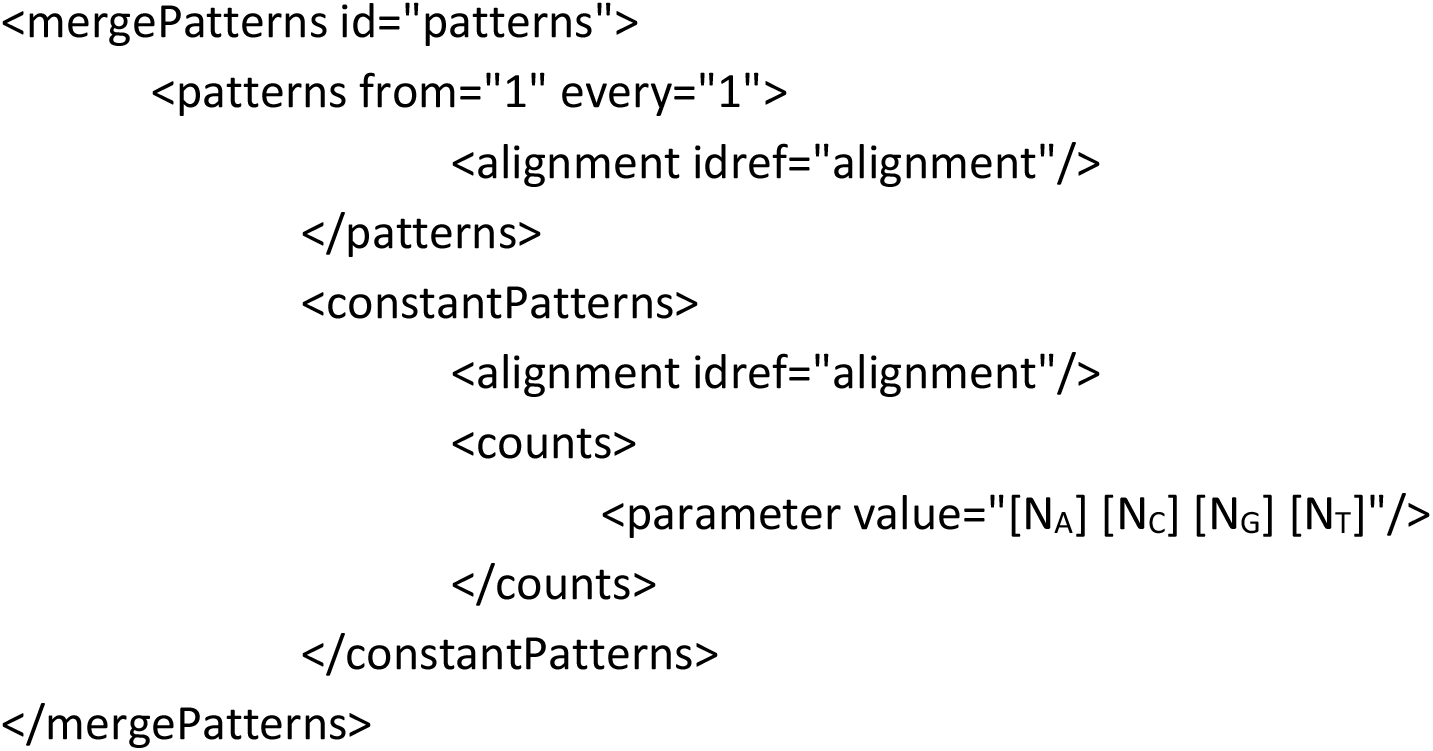

The resulting modified XML files were then used for runs of BEAST v1.10.4 with 100,000,000 MCMC iterations. We observed rapid convergence of the associated posterior log-likelihood traces. The first 10,000,000 iterations (10%), were consequently treated as burn-in, and maximum clade consensus trees were generated from the remaining 90% of draws using TreeAnnotator v1.10.4 (*27*). The resulting maximum clade consensus trees and highest posterior density intervals were displayed using *ggtree* v3.2.0 (*82*).

##### Estimating DFT1 and DFT2 origin dates

There are three levels of uncertainty in estimates of DFT1 and DFT2 dates of origin. First, there is uncertainty in mutation rate (Table 2). Second, there is uncertainty as to the germline or somatic classification of variants occurring at the trunk of the DFT1 and DFT2 phylogenetic trees. Such variants fulfil the technical definition of “somatic” (i.e. presence in ≥1 tumours and absence in all 80 normal devil genomes); however, it is not possible to determine the exact proportion of such variants which are, in fact, rare germline variants that occurred in the DFT1 or DFT2 founder devils. We identified 1,311 DFT1 truncal variants and 1,335 DFT2 truncal variants. Third, there is uncertainty regarding the date represented by the clonal mutations obtained from a polyclonal tumour biopsy (see section “Tumour heterogeneity and tip dating” below).

In order to obtain a best estimate of the proportion of truncal variants which are germline, we counted the number of variants unique to each normal Tasmanian devil genome. Among 58 devils of “Eastern” ancestry (Figure S2), there was a median of 818 (range 237-1524) singleton substitution variants. We thus designated the most likely breakdown of truncal variants in DFT1 and DFT2 as follows:

- DFT1: 818 germline variants, 493 somatic mutations (range of germline variants, 0 – 1,311)
- DFT2: 818 germline variants, 517 somatic mutations (range of germline variants, 0 – 1,335)

The origin dates for DFT1 and DFT2 were estimated by applying the mean clock rate estimates provided by BEAST (Table 2) to the respective estimated number of truncal somatic mutations in DFT1 and DFT2 in order to obtain time intervals. These time intervals were then subtracted from the mean most recent common ancestor (MRCA) dates outputted by BEAST, resulting in the date-of-origin estimates of 1986.435 (June 1986) for DFT1 and 2011.237 (March 2011) for DFT2.

The upper and lower origin date boundaries were derived as follows. We first extracted the parameter values outputted in 90,000 MCMC draws after burn-in (1 every 1,000 iterations) obtained from runs of BEAST v1.10.4. The upper boundary was set as the 95^th^ percentile of the highest posterior density distributions for MRCA date; this assumes that the entirety of truncal variants are germline and that MRCA date is equivalent to the origin date. These estimates were 1989.039 (January 1989) for DFT1 and 2012.346 (May 2012) for DFT2. For the lower boundary, we applied the corresponding mutation rate to the entirety of truncal mutations, with the assumption that all are somatic, and subtracted this from the corresponding MRCA date. We obtained the 5^th^ percentile of the highest posterior density distributions around the lower origin date, which resulted in lower boundary origin dates of 1982.264 (April 1982) for DFT1 and 2009.508 (July 2009) for DFT2.

##### Tumour heterogeneity and tip dating

We note that the bulk tumour sequencing approach used here generally only allows identification of clonal variants present within all cells within the sampled biopsy. Thus, each tumour sequence reported here represents that of the most recent common ancestor of the cells within the sampled tissue. In DFT, the most recent common ancestor usually represents the dominant cell transmitted at infection, which probably in most cases occurred months or even years prior to tumour sampling. However, for simplicity, and due to lack of knowledge of this offset, we assigned tip dates as sampling dates. It should be acknowledged therefore that, for this reason, the dates of tips, nodes and ancestors in the phylogenetic trees presented here are likely positively offset by a period of several months (e.g. a tumour sampled in July 2015, and labelled as occurring on this date, may, for instance, represent a cell that occurred in July 2014). This phenomenon, known as the pretransmission interval, has been recognised in molecular phylogenetics (*51*) and tumour coalescent theory (*85*).

#### 4.7 Intra-tumour heterogeneity analyses

We screened DFT1 and DFT2 tumour genomes for the presence of subclones by visually inspecting somatic substitution variant allele fraction (VAF) distributions and searching for populations of variants with lower than expected VAF. We computationally isolated candidate subclones using VAF thresholds on variants subsetted by tumour sharing patterns. Maximum likelihood phylogenetic trees were then generated with these sequences using RAxML-NG v1.0.3 (*81*). One hundred starting trees steps were used in combination with model parameters GTR+G+ASC_LEWIS. Highest likelihood trees were displayed using *ggtree* v3.2.0 (*82*).

#### 4.8 Identification of recurrent substitutions and indels

BEAST maximum clade consensus trees for DFT1 and DFT2 were processed using the *phangorn* v2.7.0 *R* library (*86*). We used the ‘allDescendants’ function to reconstruct all possible tip and internal branch ancestral states within the DFT1 and DFT2 lineages, and screened for substitutions and indels which were discordant with the tree topology. We visually validated candidate recurrent mutations involving protein-coding gene exons using IGV (*87*). Validated recurrent variants are annotated in Table S7 (in DFT1 affecting *ECEL1*, *ENSSHAG00000018547;* in DFT2 affecting *NT5EL*).

#### 4.9 Variant gene annotation

Substitutions and indels were annotated using the Ensembl Variant Effect Predictor (*88*) with default settings and mSarHar1.11 Ensembl v104 gene annotation. We flagged genes included in the COSMIC v.94 cancer gene census (*89*).

#### 4.10 Non-synonymous-to-synonymous mutation ratios

We collected DFT1 and DFT2 somatic substitution and indel mutations occurring within protein-coding gene exons or at essential splice sites (see section 4.9). Mutations exclusively found among six tumours with <70% purity and lacking sequences from matched hosts (DFT1: 208T2, 837T1a, 1071T1, 2692T; DFT2: 1553T1, 1553T2) were excluded. Recurrent variants (see section 4.8) were supplied as multiple entries. Global dN/dS ratios were calculated using *dNdScv* v0.1.0 (*33*) with a custom devil reference coding sequence (’RefCDS’) database built from Ensembl v104 gene annotations of mSarHar1.11 (19,005 protein coding genes on chromosomes 1-6, X and Y), specifying flags cv = NULL, max_muts_per_gene_per_sample = 100, max_coding_muts_per_sample = 10000, min_indels = 1.

### 5. Rearrangements

#### 5.1 Rearrangement calling and genotyping

Somatic rearrangements were called and genotyped in DFT1 and DFT2 using two software packages, SvABA (*90*) and MSG (https://github.com/MaximilianStammnitz/MSG).

##### SvABA

Structural variants were joint-called across tumours (-t; DFT1 tumours, DFT2 tumours and the single non-transmissible carcinoma), as well as normal (-n) devil samples using SvABA v1.1.3 (*90*). The output file svaba.somatic.sv.vcf.gz was processed as follows. We first restricted consideration to candidate rearrangements involving chromosomes 1-6, X, Y and MT, and with QUAL ≥25/100. We then retained only rearrangements that: (1) occurred with ≥6 reads in at least one tumour with purity ≥20% and (2) showed no evidence of germline presence, defined as occurrence with ≥3 reads in one of more of the 80 normal devil genomes or presence with ≥6 reads across all normal genomes combined.

Inspection of remaining candidate rearrangements revealed that 5-10% of these grouped in closely related clusters, and appeared to be artefacts produced by a failure to merge duplicate calls. We identified such clusters by searching for candidate rearrangements with close coordinates (both left and right breakpoint coordinates <500 bp from one another), and identical strand and genotype predictions. For each cluster, we retained only the entry with the highest QUAL value or, in cases where entries had identical QUAL value, the highest total read support across tumours.

We used hierarchical clustering of rearrangements to assess phylogenetic relationships among tumours, and compared this with phylogenetic trees produced using substitution variants (see section 4.6). Concordance was maximised with “presence” of a rearrangement defined as ≥3 supporting reads. This definition was therefore used to define rearrangement presence and absence among tumours.

##### MSG

Rearrangements were called in each individual tumour and normal BAM file using Manta v1.6.0 (*91*), followed by reformatting of inversions (https://github.com/Illumina/manta/blob/master/src/python/libexec/convertInversion.py) in the “tumorSV.vcf” output. Next, we used svimmer v0.1 (*92*) to merge calls from all samples, using a maximum breakpoint distance of 500 bp on both ends of structural variants as a threshold for merging. We then genotyped the final input list of rearrangements across all samples using GraphTyper v2.6.1 (*92*).

The resulting genotyping matrix was processed as follows. Only hits involving chromosomes 1-6, X, Y and MT were retained. Using the *vcfR* v1.12.0 library in *R* (*93*), variant allele fractions (VAFs) of structural variants were calculated using the allelic depth (’AD’) information field. We then removed candidate rearrangements with >1 alternative (ALT) genotype, with QUAL=0, or which showed 100% REF or 100% ALT read support across all samples. For deletions, VAFs were determined using Graphtyper’s breakpoint genotyping model (’DG.0’), whereas VAFs of inter-chromosomal translocations (’OG’) were calculated using only reads derived from the left-hand or right-hand breakpoint, whichever had the higher QUAL value.

Next, we retained only rearrangements that: (1) occurred with ≥6 reads in at least one tumour with purity ≥20%; (2) showed no evidence of germline presence, defined as occurrence with ≥3 reads in one of more of the 80 normal devil genomes or presence with ≥6 reads across all normal genomes combined; and (3) showed a median total coverage of ≥20 reads (both REF and ALT) in tumours.

We inspected remaining rearrangements for clusters of similar rearrangements, which may come about due to failure to merge duplicate calls. We identified such clusters by searching for candidate rearrangements with close coordinates (both left and right breakpoint coordinates <500 bp from one another), and identical strand and genotype predictions. Three such clusters were identified, and in each cluster, we retained only the variant with highest QUAL value.

We used hierarchical clustering of rearrangements to assess phylogenetic relationships among tumours, and compared this with phylogenetic trees produced using substitution variants. Concordance was maximised with “presence” of a rearrangement defined as ≥1 supporting reads. This definition was therefore used to define rearrangement presence and absence among tumours.

#### 5.2 Merging rearrangements called with SvABA and MSG

We merged SvABA and MSG rearrangements if both their left-hand and right-hand breakpoint coordinates overlapped within <500 bp, and if both breakpoint strand orientations were also identical.

#### 5.3 Rearrangement breakpoint assembly

Subsequent to rearrangement set merging, we re-assembled rearrangement breakpoints to base-pair resolution using TIGRA-SV v0.4.3 (*94*). The algorithm was supplied with input coordinates from MSG for structural variants supported by both MSG and SvABA. TIGRA-SV parameters used were -l = 1000 [maximum bp from breakpoint], -a = 100 [maximum bp into breakpoint], -w = 400 [bp of additional reference padding on both ends], -p = 50000 [max. depth], -h = 5000 [maximum graph nodes in assembly].

83% DFT1 and 90% DFT2 rearrangements could be re-assembled by TIGRA-SV. For these, initial reference breakpoint coordinate positions were occasionally updated by slightly offset coordinates supported by output from TIGRA-SV. This led us to identify and remove a small number of additional rearrangements which likely represented duplicate entries. Entries were defined as potential duplicates if, in both cases both left-hand and right-hand coordinates mapped <500 bp from one another, and if identical strand and genotypes were predicted. In such cases we retained only the entry with highest QUAL value or, in cases where entries had identical QUAL value, the highest total read support across tumours.

Rearrangements were defined as somatic LINE-1 insertions if at least one breakpoint reassembly indicated untemplated poly-A or poly-T of ≥10 bp. These were excluded from rearrangement analysis, and instead included in transposable element analysis (see section 6.3).

These steps produced a final genotyped set of 2,868 somatic rearrangements (2,120 in DFT1; 533 in DFT2; 215 in the non-transmissible carcinoma) (Table S5).

#### 5.4 Rearrangement event clustering

We collapsed rearrangements into “rearrangement events” defined as groups of physically connected rearrangements co-occurring in phylogenetically concordant subsets of tumours.

Substitution phylogenetic trees for DFT1 and DFT2 (see section 4.6) were processed using the *phangorn* v2.7.0 *R* library (*86*). We used the ‘allDescendants’ function to reconstruct all possible tip and internal branch ancestral states among tumours. Intersections with rearrangement calls revealed 83 DFT1- and 37 DFT2-specific nodes (ancestral, internal or tip) supported by at least two MSG-called rearrangements. A network graph was generated for each of these nodes, in which vertices represent rearrangements and weighted edges represent the minimum genomic distance between breakpoints of two proximal rearrangements. An upper distance threshold of 500 kb was employed for linking breakpoints along the same chromosome. Within each phylogenetic node, we then identified and merged event groups of spatially interlinked rearrangements by running 100 iterations of the Markov clustering (MCL) (*95*) algorithm on the associated edge matrix, setting parameters to ‘expansion’ = 4 and ‘inflation’ = 1, 2, or 3.

This process merged 1,589 rearrangements into 1,385 rearrangement events in DFT1, and 402 rearrangements into 342 rearrangement events in DFT2. For the non-transmissible carcinoma, 340T, we included both MSG and SvABA rearrangements in the clustering approach, merging 215 rearrangements into 60 rearrangement events.

#### 5.5 Rearrangement event rates

Rearrangement events assigned to each tumour were counted, and counts were regressed against sampling date using *R* (*84*), excluding tumours for which exact sampling date was unknown. 95% confidence intervals and R^2^ values were calculated using the ‘lm’ function in R (*84*).

#### 5.6 Rearrangement gene annotation

Rearrangement breakpoint coordinates were annotated against both Ensembl (mSarHar1.11, v104) and RefSeq (mSarHar1.11, v103) gene coordinates. We additionally flagged any breakpoints falling into genes listed in the COSMIC v.94 Cancer Gene Census (*89*) (Table S5).

### 6. LINE-1 retrotransposition

#### 6.1 LINE-1 annotation in mSarHar1.11

We annotated LINE-1 elements in mSarHar1.11 using RepeatMasker v4.0.8 in sensitive mode (*96*) with the following settings: RepeatMasker -engine wublast -species ‘sarcophilus harrisii’ -s -no_is -cutoff 255 -frag 20000. This identified 932,788 LINE-1 element hits (537.8 Mbp, covering 17.4% of the total mSarHar1.11 genome). 1,948 of these likely correspond to full-length LINE-1 elements >6.3 kb (Table S1).

#### 6.2 LINE-1 insertion calling and genotyping

Two tumours, 18T and 1538T1, were excluded from LINE-1 analysis for technical reasons (low DNA quality and short library insert size, respectively). We used RetroSeq v1.5 (*97*) to identify LINE-1 insertions in DFT1 and DFT2 tumours. We first curated a full-length LINE-1 element (Chr1: 516,591,954 (3’) - 516,598,560 (5’)) for use in realignment steps. For each input BAM file, the discordant read discovery step was run as follows: retroseq.pl -discover -bam input.bam -output out.reads -eref L1_ref_path.txt -align; and the LINE-1 calling step was run as follows: retroseq.pl -call -bam input.bam -input out.reads -ref mSarHar.fa -output out.vcf -reads 10 -soft.

We filtered each of the resulting RetroSeq VCF files to exclude entries in which both the left-hand and right-hand breakpoints failed the discordant read ratio test (filtering level <6 out of 8). Next, we merged entries from all individual VCF files by dividing the genome into non-overlapping windows ≥1 kb and combining entries that fell within the same bin. This yielded 125,220 windows with evidence of LINE-1 insertions. We used this to generate a presence/absence matrix by genotyping each of the 125,220 windows in each sample. “Presence” was defined as passing RetroSeq filtering level ≥6 out of 8. After removing any windows that showed presence in ≥1 normal genomes, we reduced the set to 7,398 windows containing putative somatic LINE-1 insertions in DFT1, DFT2 or the non-transmissible devil carcinoma. Remaining candidates were additionally filtered to retain only those for which ≥1 tumours passed all RetroSeq filters (filtering level 8 out of 8). This produced a final set of 937 candidate somatic LINE-1 insertions (120 in DFT1, 813 in DFT2, 4 in the non-transmissible carcinoma 340T).

#### 6.3 Merging LINE-1 insertion called with RetroSeq, SvABA and MSG

We integrated putative LINE-1 mobilisations from the SvABA and MSG rearrangement sets (see section 5) into the list of somatic LINE-1 insertions (Table S4). SvABA and MSG rearrangement calls were defined as LINE-1 insertions if at least one breakpoint reassembly output from Manta or TIGRA-SV indicated untemplated poly-A or poly-T tracts of ≥10 bp. This produced a final set of 1,141 candidate somatic LINE-1 insertions (135 in DFT1, 981 in DFT2, 25 in the non-transmissible carcinoma) (Table S4).

#### 6.4 LINE-1 3’ transduction analysis

We searched for LINE-1 3’ transductions in DFT1 and DFT2 by screening the candidate LINE-1 insertions (called by RetroSeq, SvABA or MSG) for those in which either the left-hand or right-hand breakpoint contained discordant reads mapping to a unique chromosomal location (representing a unique genomic region 3’ to a LINE-1 source element). First, chromosomal regions of mSarHar1.11 (excluding unmapped scaffolds and MT) were subdivided into 20 kb windows. For each candidate LINE-1 insertion, discordantly mapped reads at each breakpoint (left and right) were collected, and the chromosomal mapping location of each discordantly-mapped mate was assigned to one of these 20 kb windows (“mate-windows”). We counted the number of reads in each mate-window, and retained only those LINE-1 breakpoint:mate-window pairs for which a single 20 kb mate-window carried three times more discordantly-mapped mates than any other mate-window. These were further filtered (1) to conservatively remove LINE-1 breakpoint:mate-window pairs whose mate-window contained <15 discordant reads associated with its corresponding candidate LINE-1 insertion and (2) to retain only those LINE-1 breakpoint:mate-window pairs for which ≥75% LINE-1 breakpoint-associated discordantly-mapped reads within the mate-window occurred on a single strand. 94 LINE-1 insertions satisfied these conditions for 3’ transduction (9 in DFT1, 83 in DFT2, 2 in 340T the non-transmissible carcinoma) (Table S4).

#### 6.5 LINE-1 insertion and 3’ transduction validation

In order to assess the accuracy of our LINE-1 insertion predictions, we randomly selected 60 LINE-1 insertion candidates (20 DFT1-specific; 20 DFT2-specific; 20 340T (non-transmissible carcinoma)-specific) for visual validation using IGV (*87*). In each case, we qualitatively assessed the likely validity of a candidate LINE-1 insertion by scoring the presence of the following hallmarks: (1) target-site duplication of 5-20 bp; (2) target site flanking reads with soft-clipped poly-A or poly-T sequences; (3) discordant reads mapping to LINE-1 elements; (4) in the case of 3’ LINE-1 transduction, a strand-biased discordant read cluster mapping to a single locus in the reference genome; (5) in the case of 3’ LINE-1 transduction, copy number amplification at the putative source locus. This screen revealed that 3/20 (14%) DFT1 candidates, 18/20 (90%) DFT2 candidates, and 19/20 (95%) 340T candidates were likely to be true somatic LINE-1 insertions.

#### 6.6 LINE-1 insertion gene annotation

We intersected LINE-1 insertion breakpoints with genes annotated by Ensembl (mSarHar1.11, v104) and NCBI (mSarHar1.11, v103) (Table S4), flagging genes included in the COSMIC v.94 cancer gene census (*89*).

### 7. Copy number variants

#### 7.1 Copy number variant calling – total copy number

##### Sequencing depth calculation

Sequencing reads were aligned to mSarHar1.11 using BWA-MEM (*78*). Read depth in 1 kb non-overlapping genomic bins, excluding the mitochondrial genome and unassigned contigs, was counted across tumour and normal genomes using bedtools v2.23.0 subcommands ‘makewindows’ and ‘coverage’ (*79*). Bins for which median read depth among normal samples was zero were discarded.

##### Tumour LogR calculation and denoising

Read counts from normal genomes were used to produce noise-corrected estimates of the log_2_ tumour:normal read depth ratio (logR) for each sample following the tangent normalisation method used in the Genome Analysis Tool Kit (GATK) (https://github.com/broadinstitute/gatk/blob/master/docs/CNV/archived/archived-CNV-methods.pdf), briefly described below.

We produced a matrix of binned read counts across autosomes from normal devil genomes (Table S2). This matrix excluded the following samples for technical reasons: 31H, 91H, 140H, 174H, 199H, 202H, 203H, 236H, 378H, 528H, 812H, 2693H). Columns and rows represented samples and bins, respectively. We normalised the matrix by dividing columns by their median value, then by dividing rows by their median value. Zero values were replaced by their row’s median. Row medians were retained for logR computation (see below). For each sample, values falling below the 0.001 or above the 0.999 quantiles of the distribution of normalised read counts were clamped to the nearest of these quantiles. Column normalisation was then repeated, and base-2 logarithm calculated for each entry of the matrix. Singular value decomposition was performed on this normalised log_2_ matrix, and left-singular vectors retained.

To calculate tumour logR values, we arranged the tumour binned read counts in a matrix, with columns corresponding to samples. These were normalised by dividing columns by their medians. The rows of the resulting matrix were then divided by the row medians derived from the normal samples. We computed base-2 logarithm of the entries in the resulting matrix, to produce a logR matrix.

Finally, the singular vectors were used to remove systematic noise from the logR matrix according to the following equation (M = logR matrix, M* = denoised logR matrix, U_K_ = matrix of K singular vectors, where K = number of singular vectors used for denoising; T denotes matrix transpose):

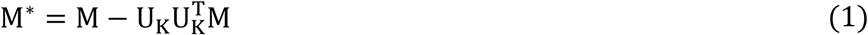

We used the ten singular vectors associated with the ten largest singular values to denoise the logR matrix.

For the sex chromosomes, binned read counts from normal genomes were segregated by sex, producing a matrix of exclusively male genomes and one of female genomes. The columns of these matrices were divided by the median depth of the equivalent autosomal genome. The remainder of the normalisation was done in the same manner as for the autosomal matrix, but the logR of the sex chromosomes of tumours derived from male or female hosts were calculated by dividing by the row medians of the host matrix of the same sex. Sex chromosomes of cell lines with 100% purity (Table S2) were normalised using the female matrix. Ten singular vectors were used for denoising.

##### Copy number transformation

For each 1 kb bin, we transformed logR (*r*) to tumour total copy number (*N_T_)* using the following equation, following a described method (*98*).

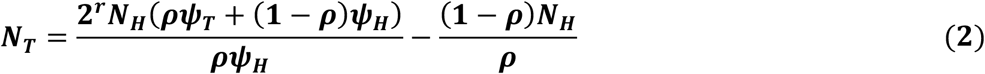

We assume that tumour ploidy (Ψ_T_) is 2 for diploid tumours and 4 for tetraploid tumours (see section 7.5 on tetraploidy), the host ploidy (Ψ_H_) is 2, and the host copy number (N_H_) is 2, except for the X and Y chromosomes of male samples, which we assume to each be at copy number 1, and the Y chromosome of female samples, which we assume to have copy number 0. The tumour purity, ρ, is estimated using the read count method described in section 3.2 (Table S2).

##### Estimation of copy number for normal samples

We used the matrix of read counts per 1 kb bin for normal genomes as used in the denoising panel. To obtain logR for the normal samples, we did the following: divide each column by its median calculated only over autosomal positions; divide each autosomal row by the previously calculated row median; divide the entries of the sex chromosome rows belonging to male or female samples by the row medians previously obtained from the corresponding sex-restricted matrix; for each sample, calculate the copy number using equation 2, assuming a ploidy (Ψ_T_) of 2 and a purity (ρ) of 1.

##### Germline copy number polymorphism

Regions of germline copy number polymorphism produce unreliable estimates of tumour copy number, as it is generally not possible to determine whether an observed apparent change in copy number is due to a tumour or a host variant, or to an artefact of the denoising correction. For this reason, any regions of the genome carrying germline copy number polymorphism were excluded from copy number segmentation and copy number estimation.

In each normal devil genome, copy number data was segmented using the Piecewise Constant Fitting (PCF) algorithm, as outlined below (see below section “Segmentation and copy number assignment” for details). The copy number of each segment was determined as follows. We first manually curated germline copy number polymorphisms on chromosome 6 in order to obtain training data for a neural network. Copy number data within each segment within the training set were converted via kernel density estimation to a vector of length 128, using the *R* function ‘density’ (*84*) applied over the range 0-5. These 128 values were used as the input layer to a feed-forward neural network with two hidden layers of sizes 32 and 8, and an output layer of size 1. The activation function used for the hidden layers was ReLU, and for the output layer, sigmoid. We used a neural network implemented using the *R torch* v0.4.0 library (*99*) to classify genomic bins as one of two categories: (1) diploid in all normal devil genomes (2) any other state. Segments for which the distribution of germline copy number was approximately normally distributed around a mean of 2 were considered to show no germline polymorphism, and were expected to produce an output of 1. All other segments were expected to give an output of 0. Training data was augmented by subsampling four data sets of at least half of the total number and at most 10,000 points from each training segment, with a fifth subsample used for validation. The neural network was trained for 50 epochs of 15 batches of 100 training data points, using a binary cross entropy loss function and Adam optimiser with a learning rate of 0.1.

The trained network was applied to germline copy number data across autosomes, and on chromosome X occurring in females. Any segment producing an output of less than 0.85 was considered to be polymorphic in germline copy number. Classification was manually validated by viewing histograms of copy number data within each germline segment identified as copy number polymorphic (Supplementary Data S2).

##### Bespoke DFT1 filter

In initial trial segmentations, four DFT1 samples (366T1, 378T1, 134T1 and 139T4) were associated with frequent small (<50 kb) copy number gains. Validation analyses revealed that these copy number gains were technical artefacts, probably due to low DNA quality. These were detected and excluded from the final copy number segmentation as follows: segments of 50 kb or less exhibiting an increase in copy number with respect to the preceding and following segments were identified. Of these, if the increase was observed uniquely in one of the identified samples, the segment was excluded. If the CNV was observed in between two and ten samples, at least one of which was an identified sample, we counted the number of independent phylogenetic events required to produce the phylogenetic pattern observed for the relevant CNV using the ACCTRAN method of parsimony ancestral reconstruction implemented in the *R* package *phangorn* v2.7.0 (*86*). We then took the ratio of the number of tumours carrying the CNV to the number of independent events implied by the grouping of the affected samples on the phylogeny; if this ratio was less than 3, the segment was excluded. Any excluded regions were excluded from final copy number segmentation across all samples.

##### Bespoke DFT2 filter

Similarly, in DFT2 we observed two samples (1515T1, 203T2) associated with frequent small (<50 kb) copy number events, this time including both gain and loss events. We employed a detection method similar to that used for DFT1, except that the upper limit of samples involved was reduced to three from ten, and excluded the resulting regions.

##### Segmentation and copy number assignment

We segmented total copy number data in each tumour genome, excluding bins that showed evidence of germline copy number polymorphism (see section “Germline copy number polymorphism” above). Each tumour’s copy number data were first winsorised to reduce the influence of outliers, using parameters tau = 1.5 and k = 10, and then segmented individually using the Piecewise Constant Fitting (PCF) dynamic programming algorithm (*98*), with a minimum segment size of 10 bins and a penalty parameter of 200. The breakpoints obtained were then filtered using a two one-sided t-tests (TOST) approach (*100*). In this approach, to test whether a candidate breakpoint represents a statistically supported step change in the copy number, we took the copy number data of the segments immediately to the left (A) and right (B) of the breakpoint and applied two t-tests: the first to assess whether the difference in the means of the copy number data in segments A and B is less than 0.25, and the second to assess whether the difference in the means is greater than -0.25. If both tests rejected the null hypothesis that the means differ by no more than 0.25 at the 0.05 significance level, then the breakpoint was retained, otherwise the breakpoint was removed. The value of 0.25 was selected such that retained breakpoints should roughly correspond to a change in copy number of at least a half-integer magnitude. t-tests were calculated in R using the t.test function, using the commands t.test(A, B, mu = 0.25, var.equal = TRUE, alternative = ‘less’) and t.test(A, B, mu = -0.25, var.equal = TRUE, alternative = ‘greater’) (*84*).

Per-sample breakpoints obtained after this first round of segmentation were merged into a ‘whole-dataset’ set of breakpoints. The requirement that no segment should be smaller than 10 bins was enforced by collecting groups of breakpoints falling within 10 bins of each other, computing the mean of these breakpoints’ coordinates (treating each sample’s breakpoint as an independent observation), and replacing these breakpoints with a single breakpoint at the mean position. This created a ‘whole-dataset-merged’ set of breakpoints. Each tumour was then resegmented using a restricted version of the PCF algorithm that can only introduce breakpoints at designated positions (https://github.com/kgori/segmentation, based on (*101*), algorithm 3), using the whole-dataset-merged breakpoints as candidates. The penalty parameter was again 200. This was followed by a second TOST filter, and then we calculated the integer copy number state for each segment by calculating the median of the copy number estimates within each segment, and rounding to the nearest integer. As a final filter, we merged adjacent segments that had the same integer copy number state, and recalculated the per segment integer copy number.

##### Ploidy estimation and copy number adjustment

Ploidy estimation for each tumour was done using a grid search approach. The grid search range was 1.87-2.13 for diploid samples, in steps of 0.005, and 3.74-4.26 for tetraploid samples, in steps of 0.01. For each Ψ_T_ value in the grid search, the copy number estimate was calculated according to equation 2. For each segment in the ‘whole-dataset-merged’ segmentation we calculate the median copy number and round it to the nearest integer. We then count the number of bins with copy number above or below the segment median. We select the Ψ_T_ value that makes these two counts approximately equal. A final round of copy number assignment was performed with new ploidy estimates (Table S6) using equation 2.

#### 7.2 Copy number variant calling – minor copy number

##### Substitution VAF correction

For tumours in the sample set for which we have sequenced a matched normal sample, we obtained a purity-corrected estimate of the variant allele fraction (VAF) in the tumour (V*_T_) by the following method. For each variant position we have observations of the read coverage corresponding to the alternative allele (C_ALT_), and the total read coverage (C_TOTAL_), in both the mixed tumour-normal sample and the pure normal sample. The VAF is obtained as C_ALT_ / C_TOTAL_, and we denote the VAF in the mixed tumour sample as V_T_, and the VAF in the normal host as V_H_. We derive an estimate of the probability (p) that a read in the mixed sample came from the tumour, by rearranging (*98*) equation S5 to give equation 3 (for simplicity we assume the host ploidy is 2). This probability is then used to adjust the observed mixed tumour-normal VAF to give an estimate of the pure tumour VAF (V*_T_) using equation 4.

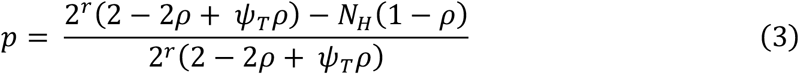

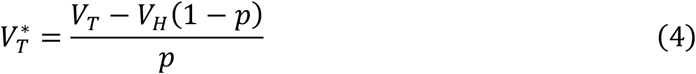

##### Minor copy number calling

We assigned a minor copy number state to each tumour sample, for each segment inferred in the copy number estimation, based on the purity-corrected germline substitution (single nucleotide polymorphism, SNP) data. This process was done in six stages: (1) SNP filtering, (2) sample selection, (3) SNP selection, (4) minor copy calling for segments with sufficient evidence, (5) minor copy calling for segments pooled together to increase shared evidence, and (6) imputation of segments with too little evidence to call. DFT1 and DFT2 clones were analysed separately.

1. *SNP filtering* We excluded from the analysis any SNPs that fell within 750 base pairs of SNPs identified by the “excessive heterozygosity filter” described in section 4.1. Additionally, SNPs occurring in regions called as copy number 1 in at least 20 tumours, but with a VAF incompatible with that copy number state (VAF >0.33 and <0.67) in more than 30% of these tumours, were excluded.
2. *Sample selection* We considered samples for minor copy calling only if they were derived from a cell line with 100% purity or had a matched normal genome available, or otherwise if the estimated purity was >0.85.
3. *SNP selection* Only SNPs that were heterozygous in the founder genotype are informative about minor copy number state. In transmissible cancer the original heterozygosity status of a SNP is unknown, but it can be inferred from the sample set by examining the VAF in each sample. Each SNP was designated as heterozygous in a sample if its VAF in that sample was between the lower and upper bounds set defined below, and its total copy number was greater than 1.

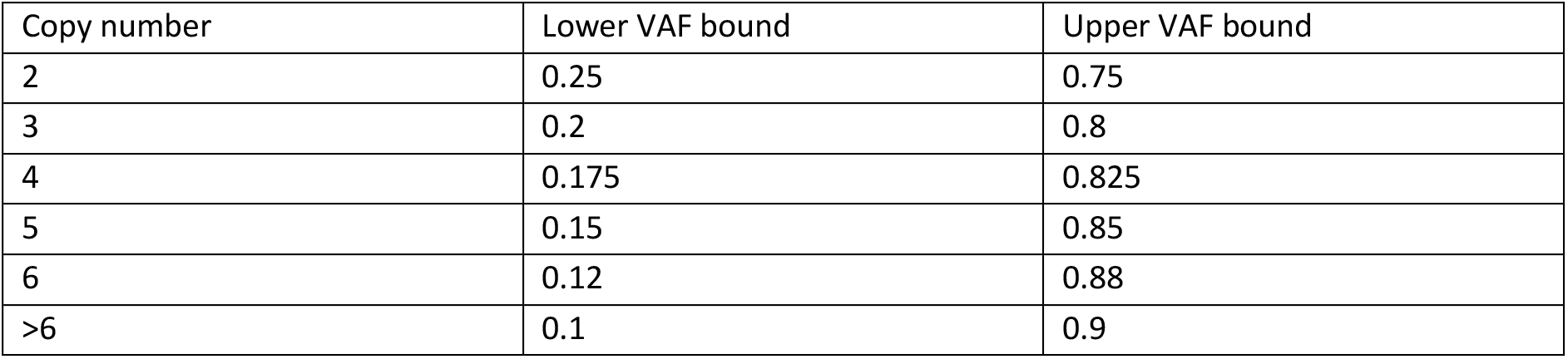 A SNP was selected as minor copy number informative if it was assigned as heterozygous in at least half of diploid tumours in which it occurred within a segment of total copy number ≥2. Subsequent minor copy number calling was done only using information from this set of informative SNPs.
4. *Minor copy calling* For each segment, we made a direct call of minor copy number if the segment contained at least 30 informative SNPs, with a minimum density of 10 SNPs per Mb. The VAF of informative SNPs was reflected in the line y = 0.5, to remove bimodality, and the mean of the reflected VAF among all SNPs in the segment was taken. Minor copy number status was assigned according to which of the intervals set out below the mean value fell into.

5. *Minor copy calling with pooling* Adjacent segments with the same assigned total copy number states in the same sample were pooled together. If the pooled segments contained at least 30 informative SNPs at a density of at least 20 per Mb, then the minor copy number state was assigned using the mean reflected VAF method used for individual segments, with the same thresholds as outlined above.
6. *Imputation* In the absence of sufficient heterozygous SNP data, minor copy state was imputed based on the minor copy number assignment of its two nearest called segments in the same sample. If these were assigned the same state, the segment was imputed to match this state. If the two nearest segments had differing minor copy calls, the segment was imputed to match the higher of the two, though for tetraploid samples, if the total copy number state was 2 and the neighbouring minor copy assignments offered a choice between 0 and 1, the minor copy number was imputed as 0. Any remaining unassigned segments occurring on autosomes were assigned a minor copy number of 1; and those on chromosome X were assigned a minor copy number of 1 in DFT1 and 0 in DFT2.

| Copy number | Minor Copy Number |  |  |  |
| --- | --- | --- | --- | --- |
|  | 0 | 1 | 2 | 3 |
| 2 | [0, 0.225) | [0.225, 0.5] | N/A | N/A |
| 3 | [0, 0.166) | [0.166, 0.5] | N/A | N/A |
| 4 | [0, 0.1) | [0.1, 0.3) | [0.3, 0.5] | N/A |
| 5 | [0, 0.1) | [0.1, 0.3) | [0.3, 0.5] | N/A |
| 6 | [0, 0.083) | [0.083, 0.25) | [0.25, 0.416) | [0.416, 0.5] |
| >6 | [0, 0.125) | [0.125, 0.5] | N/A | N/A |

#### 7.3 Copy number event reconstruction

Copy number profiles of each segment were manually inspected across the set of 78 DFT1 and 41 DFT2 tumours, and errors in copy number assignment corrected. Copy number variants (CNVs) were classified as step changes (gain of one copy, loss of one copy) from the ancestral state (Table S6). CNVs which were incongruous with the DFT1 or DFT2 phylogenetic trees were visually validated, and explained through one of the following models (1) secondary CNV interrupting a shared pre-existing CNV; (2) backmutation; (3) recurrent gain at a single locus. Such inferred events were recorded as step changes (Table S6), and phylogenetic incongruity resolved. We used MEDICC2 (*102*) to validate ancestral orders of reconstructed copy number change instances.

#### 7.4 Whole genome doubling

We assigned DFT1 and DFT2 tumours as diploid or tetraploid using two lines of evidence. (1) Tumours were defined as candidate tetraploid if a genome-wide frequency distribution of purity-corrected somatic variant allele fraction (VAF) revealed the presence of a visually detectable peak at 25%. (2) Tumours were defined as tetraploid if they carried ≥3 copy number variants (CNVs) at half-integer copy number, after having been initially assigned copy number under an assumption of diploidy (see above). This method identified 16 tetraploid DFT1s (with 15 whole genome duplication events) and 3 tetraploid DFT2 tumours (with 3 whole genome duplication events).

Timing of whole genome duplication (WGD) events was performed as follows. We used VAF thresholds appropriate for copy number state to isolate and count mutations occurring after WGD (Table S6) (*103*). After accounting for increased mutation opportunity after WGD, we applied somatic substitution mutation rates estimated using BEAST (Table 2, see section 4.6) to the post-WGD mutation counts in order to estimate a post-WGD time interval for each tumour. Uncertainty intervals were calculated using Bayesian credible intervals around the mutation rate estimate (Table S6).

#### 7.5 Copy number variant gene annotation

CNV coordinates were intersected with Ensembl (mSarHar1.11, v104) and NCBI (mSarHar1.11, v103) gene annotation (Table S6). Genes listed in the COSMIC v94 Cancer Gene Census (*89*) were additionally flagged.

### 8. Supplementary figures

**Fig. S1.**
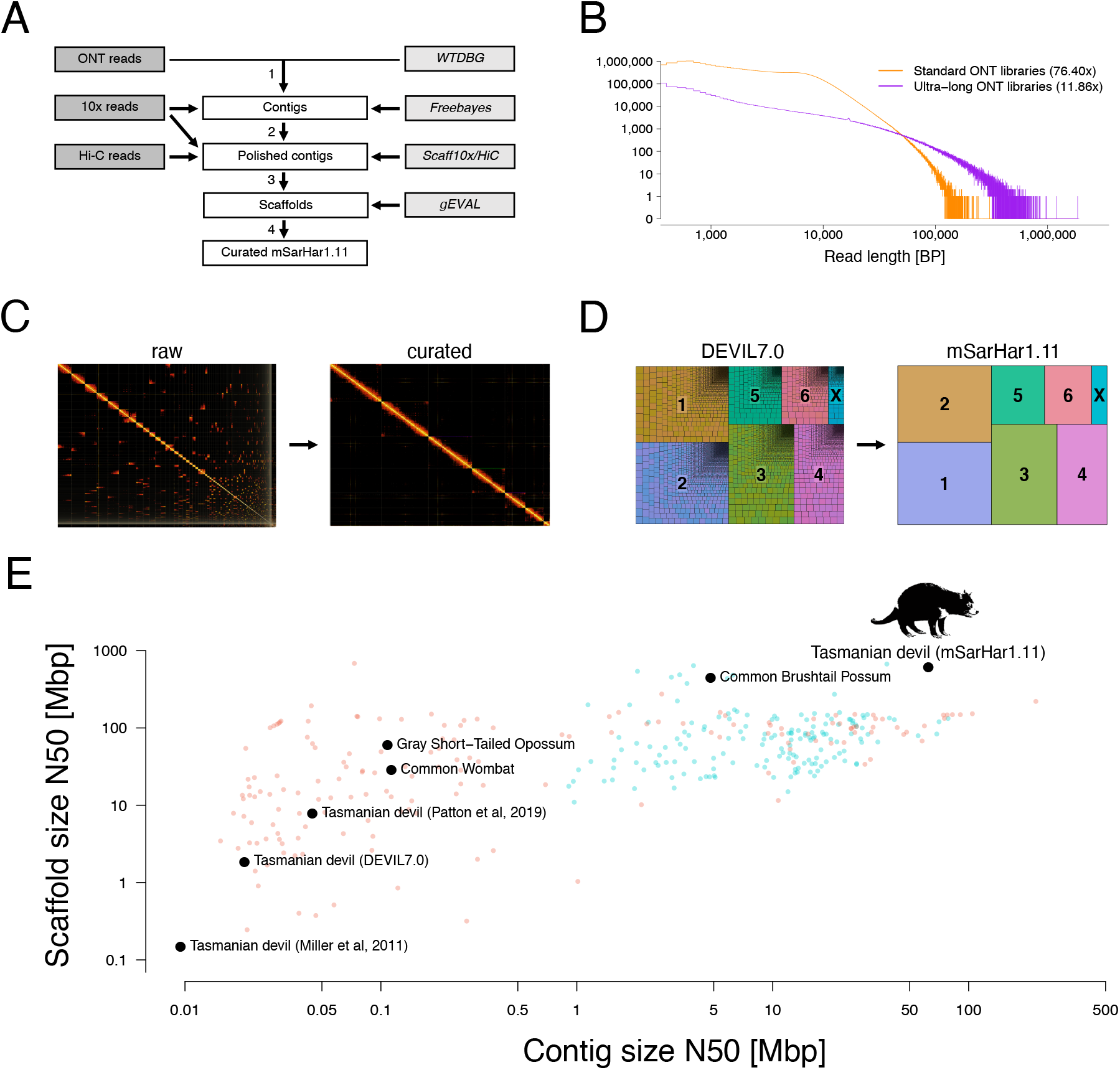
mSarHar1.11 genome assembly. (**A**) Overview of the mSarHar1.11 assembly process. (**B**) Size distribution of read lengths (base pairs) produced by standard and ultra-long Oxford Nanopore Technologies (ONT) libraries. (**C**) Hi-C chromosome-level contact map before (left) and after (right) curation. (**D**) Comparison of number of scaffolds in DEVIL7.0 (*13*) and in mSarHar1.11. Each scaffold is represented by a box, with chromosomes (1-6 and X) represented with colours. In mSarHar1.11, each chromosome is represented by a single scaffold. (**E**) Assembly statistics for various vertebrate genome assemblies (orange: NCBI RefSeq mammalian genomes; blue: VGP draft and curated genomes (*104*); summarised in Table S1). Selected marsupial genome assemblies, including previous Tasmanian devil genome assemblies, are labelled.

**Fig. S2.**
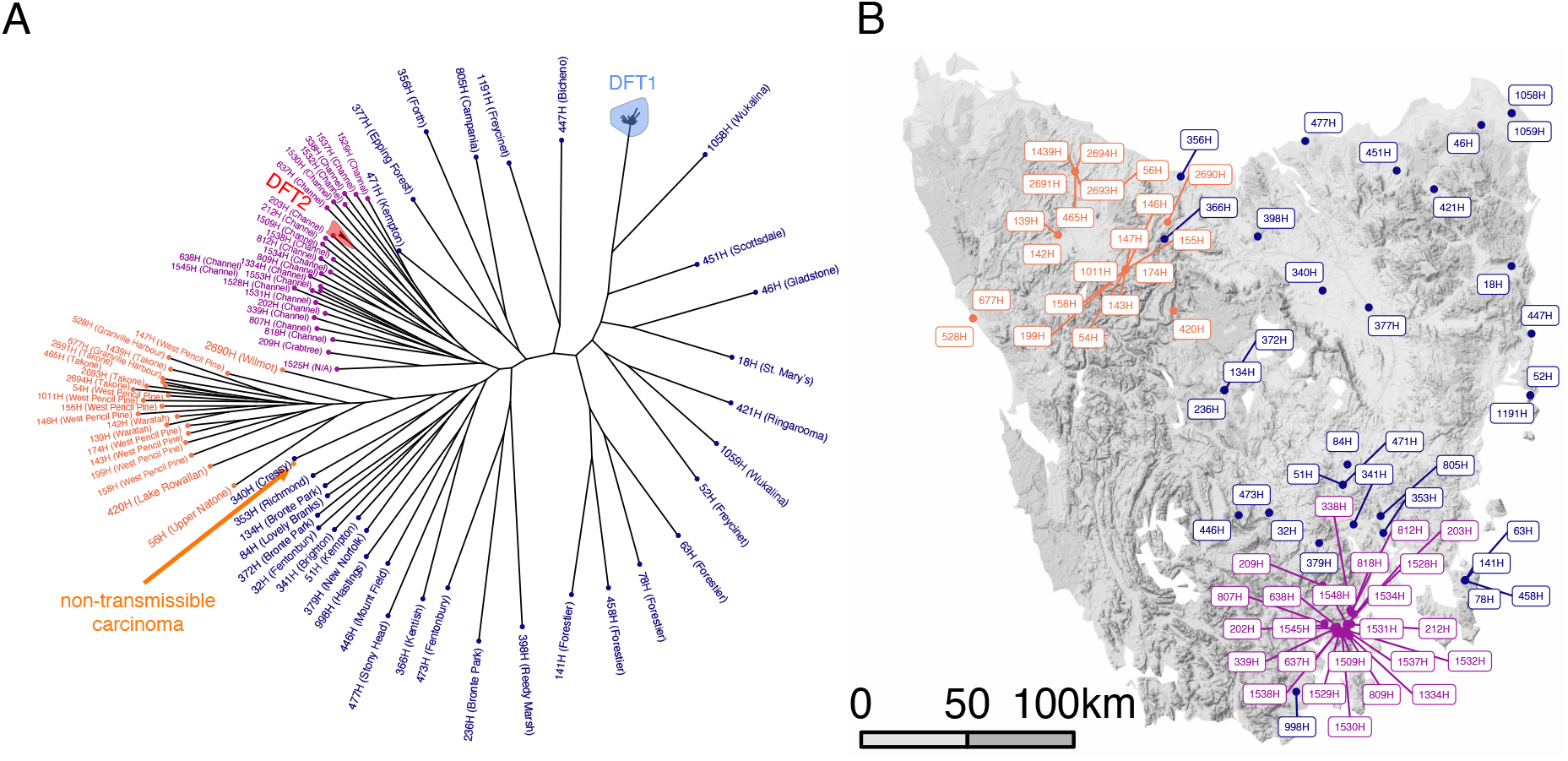
Phylogenetic tree of Tasmanian devil tumour and normal genomes. High resolution labelled version of tree displayed in Figure 1C. Normal devil genome IDs and sampling locations are labelled on tree (**A**), and corresponding locations labelled on the map (**B**), coloured by phylogenetic group (magenta, D’Entrecasteaux Channel; dark blue, Eastern Tasmania; orange, Western Tasmania). Branch lengths on phylogenetic tree are uninformative.

**Fig. S3.**
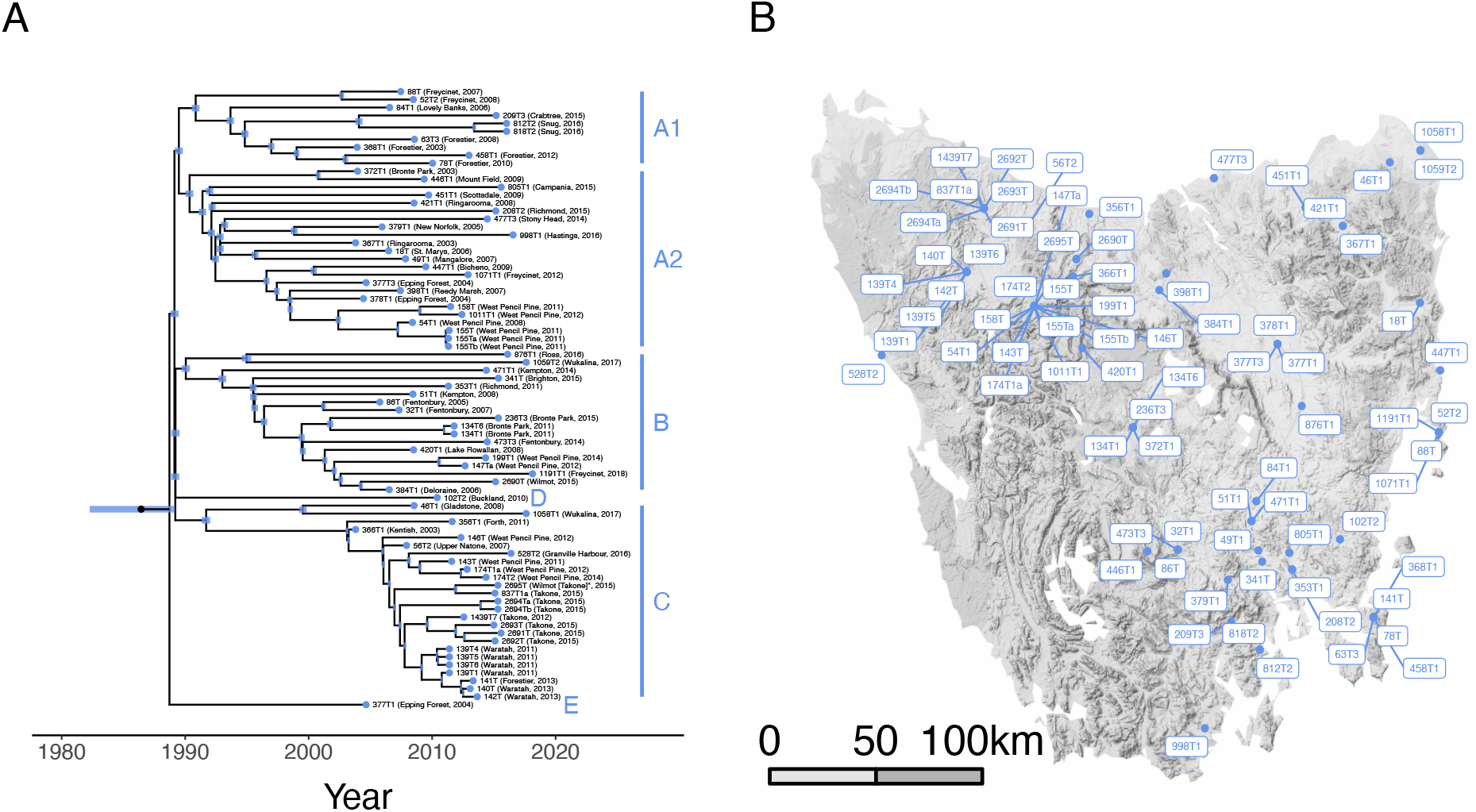
Phylogenetic tree and sampling locations of DFT1 tumours. High resolution labelled version of tree displayed in Figure 1D. DFT1 tumour IDs, sampling locations, and years of sampling are labelled on the tree (**A**), and corresponding locations labelled on the map (**B**). Tumour 2695T was reported to have been sampled in Wilmot, but its phylogenetic position, and the fact that it shares the same host as tumour 837T1a, suggest that sampling in fact occurred in Takone (asterisk).

**Fig. S4.**
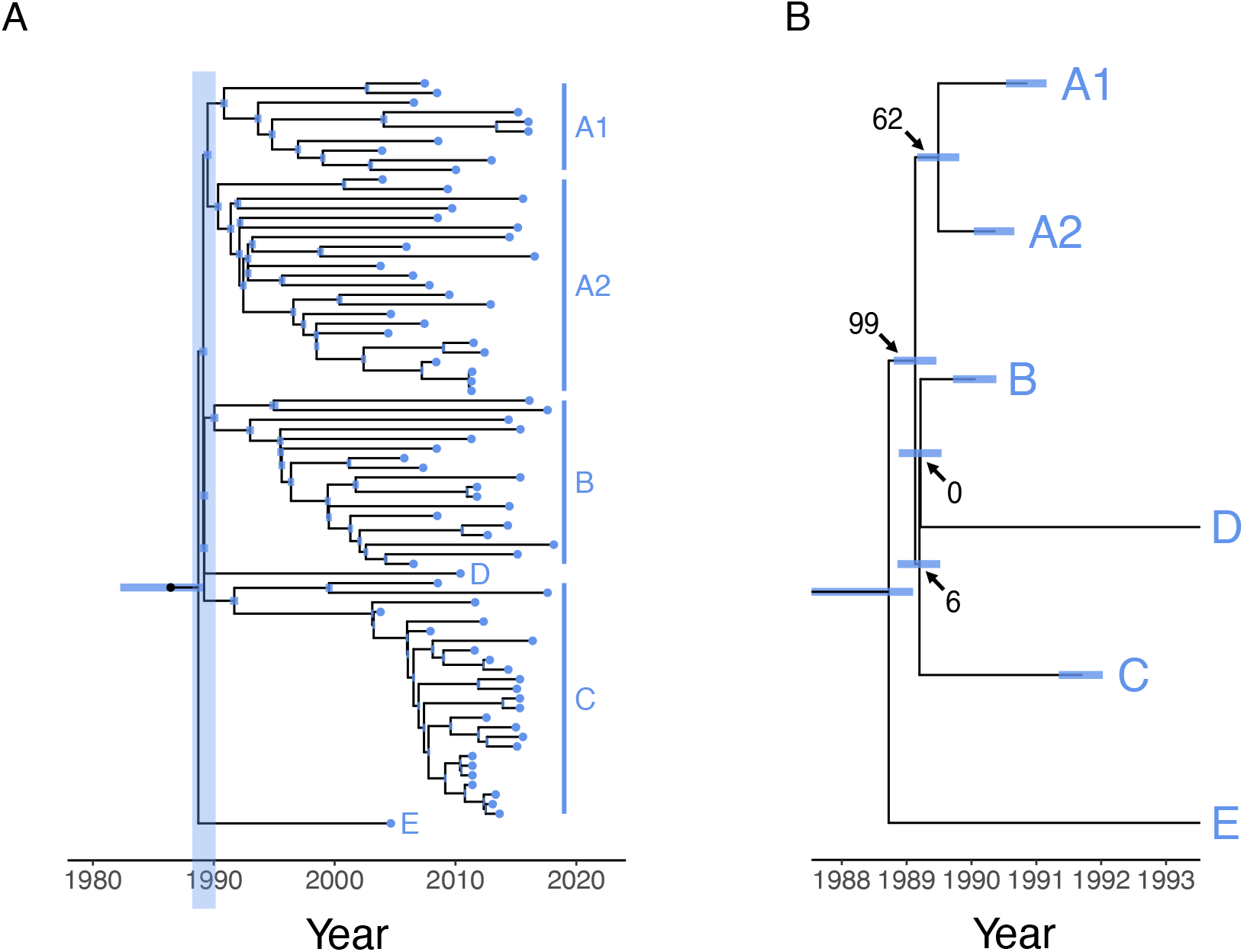
Variant counts at the base of the DFT1 phylogenetic tree. (**A**) DFT1 phylogenetic tree with “base” region highlighted. (**B**) Numbers of substitution variants at internal nodes at the base of the DFT1 phylogenetic tree. Counts are consistent with all six DFT1 clades (A1, A2, B, C, D, E) being founded with bites from a single progenitor devil in a “superspreader” event.

**Fig. S5.**
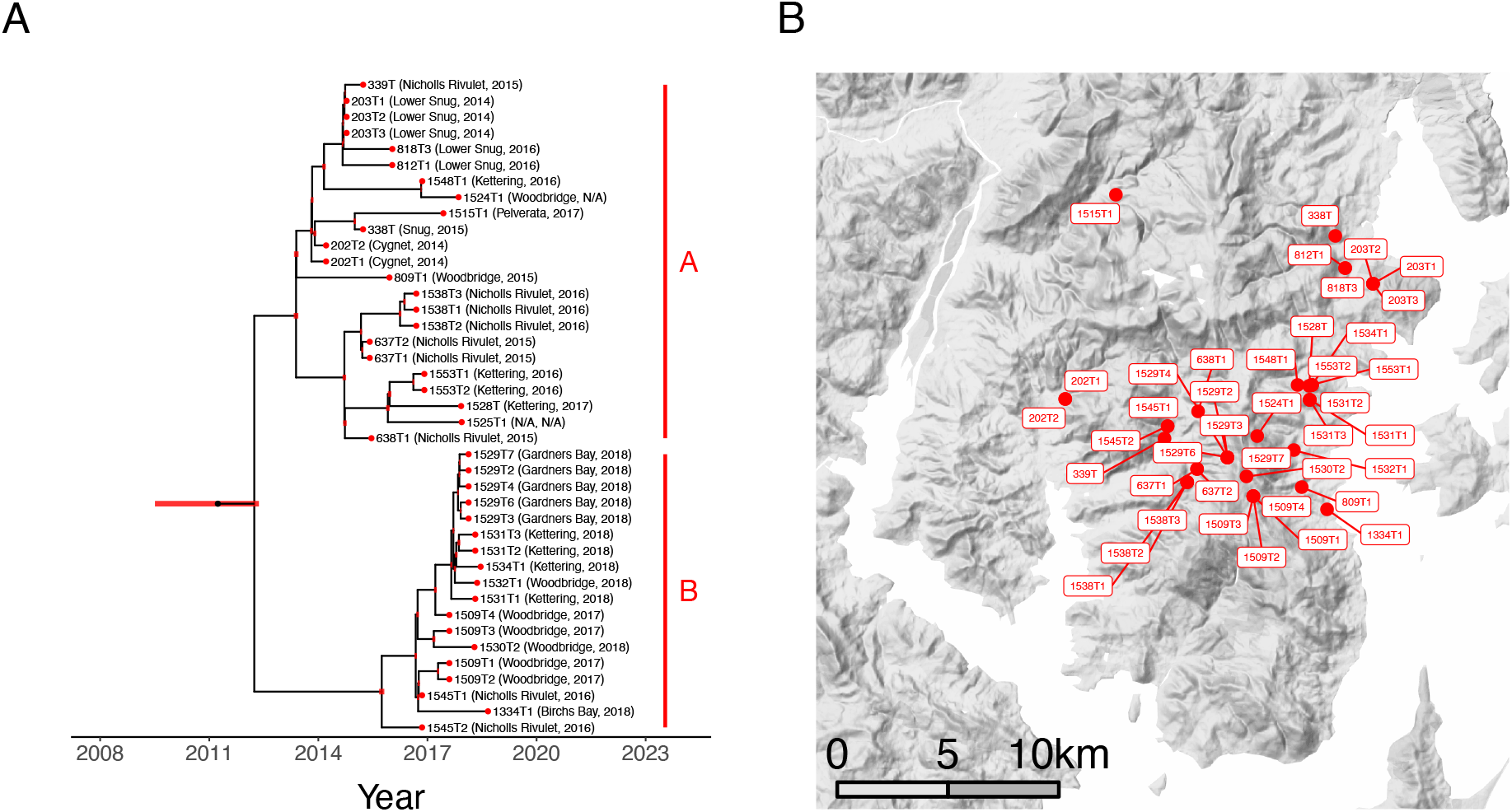
Phylogenetic tree and sampling locations of DFT2 tumours. High resolution labelled version of tree displayed in Figure 1E. DFT2 tumour IDs, sampling locations, and years of sampling are labelled on the tree (**A**), and corresponding locations labelled on the map (**B**).

**Fig. S6.**
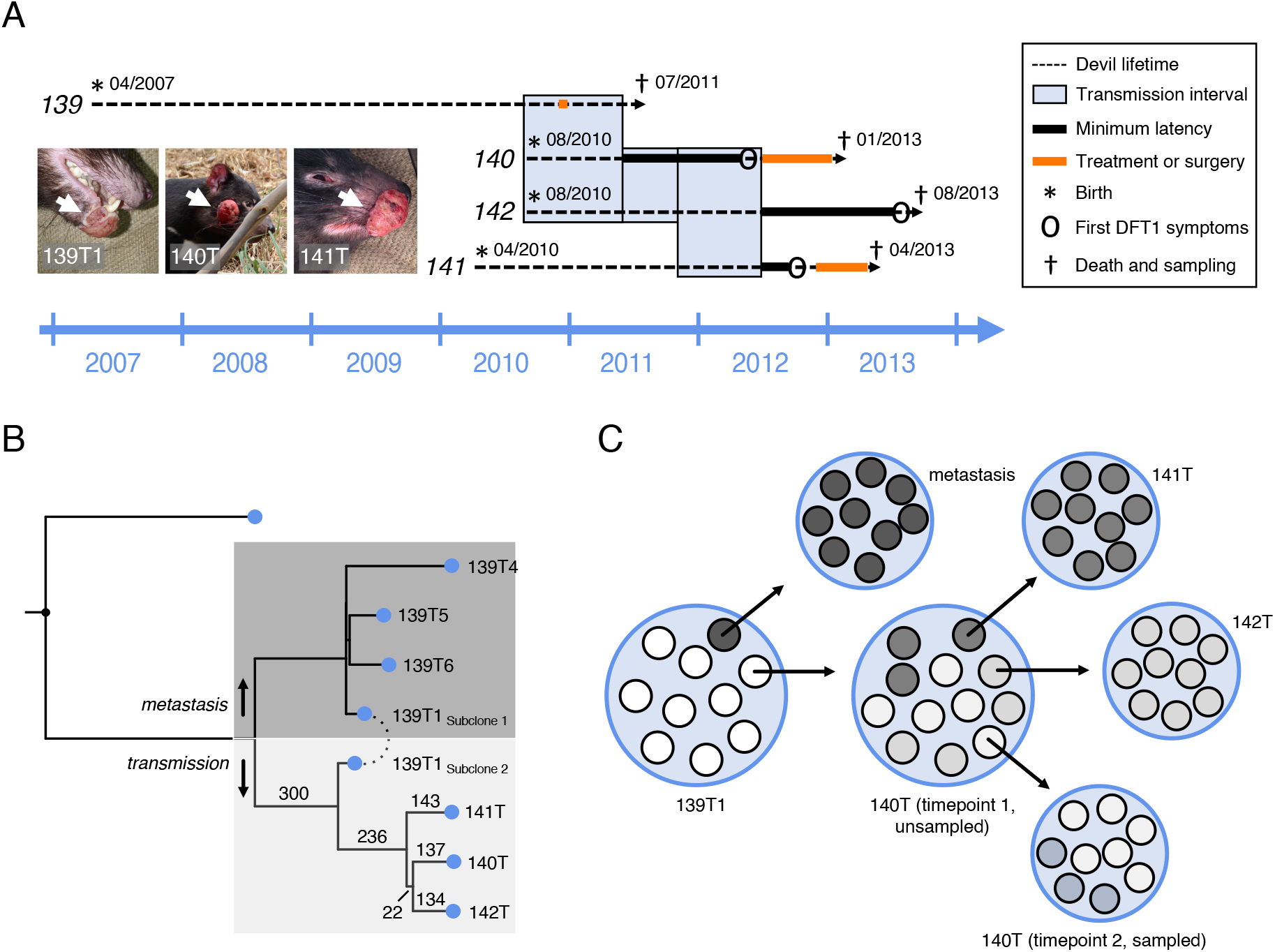
Vertical transmission of DFT1. (**A**) Detailed timeline of a DFT1 vertical transmission event, related to Figure 2. DFT1-infected female devil 139 was trapped with pouch young near Waratah in November 2010. She was subsequently housed in a care facility, together with pouch young, devils 140 and 142. Devils 140 and 142 were subsequently separated from devil 139 and housed together with another animal, devil 141, in a wildlife park enclosure. Surgical removal was attempted on devil 139’s facial tumour, and immunotherapy was attempted on devils 140 and 141. Inset photographs of sequenced tumours 139T1 (lower lip), 140T (cheek) and 141T (upper lip) support a DFT1 transmission pattern from biter (139) to bitten (140) to biter (141). While no photograph was available for tumour 142T, animal 142’s pathology report states a “pedunculated tumour at commissure of mouth, left-hand side”, and is thus suggestive of a transmission to a biter. (**B**) Phylogenetic tree illustrated in Figure 2D, with numbers of substitutions defining each branch (within the transmission cluster) marked. 22 mutations uniquely shared between tumours 140T and 142T are clonally fixed in 142T, but occur in a subclone at ∼72% frequency in tumour 140T. All 137 substitutions unique to tumour 140T were subclonal (∼67% frequency). (**C**) Model representation of DFT1 propagation between the four tumours 139T1, 140T, 141T and 142T. At least two distinctive cell populations, 139T1_Subclone1_ (dark shading, ∼10%)) and 139T1_Subclone2_ (light shading, ∼90%) were present within the tissue biopsy sampled from animal 139’s facial tumour, 139T1. Cells belonging to 139T1_Subclone2_ were transmitted to devil 139’s pouch young, devil 140, through a vertical transmission event. “140T timepoint 1” represents a model of the cellular contribution of tumour 140T at the time of onward transmission to devils 141 and 142. The 140T biopsy sequenced in this study (140T timepoint 2) was collected after several months of immunotherapy.

**Fig. S7.**
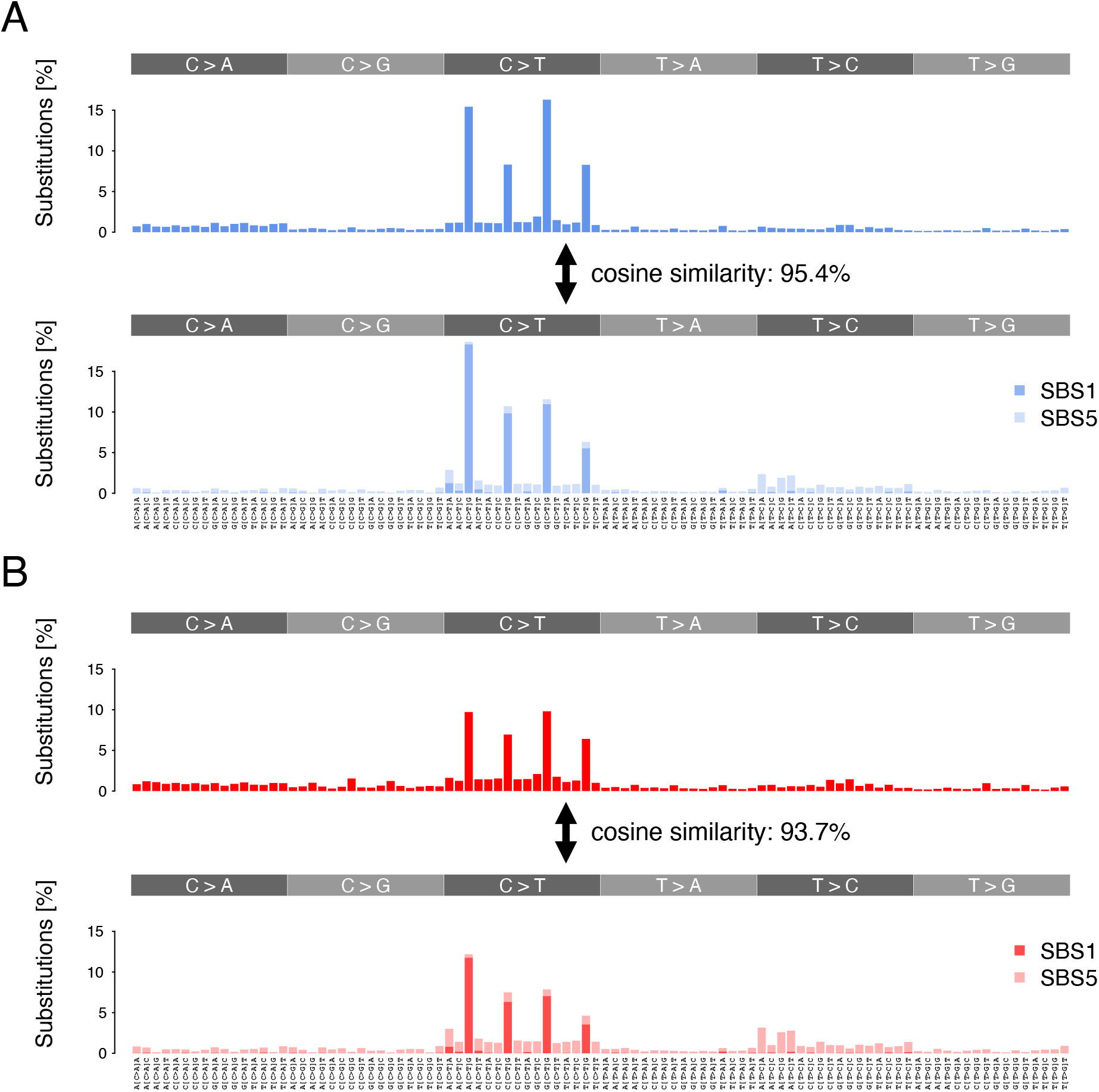
Substitution mutational spectra in DFT1 and DFT2. High resolution labelled version of DFT1 (**A**) and DFT2 (**B**) plots presented in Figure 3A. Cosine similarities with spectra reconstructed with COSMIC signatures SBS1 and SBS5 are shown.

**Fig. S8.**
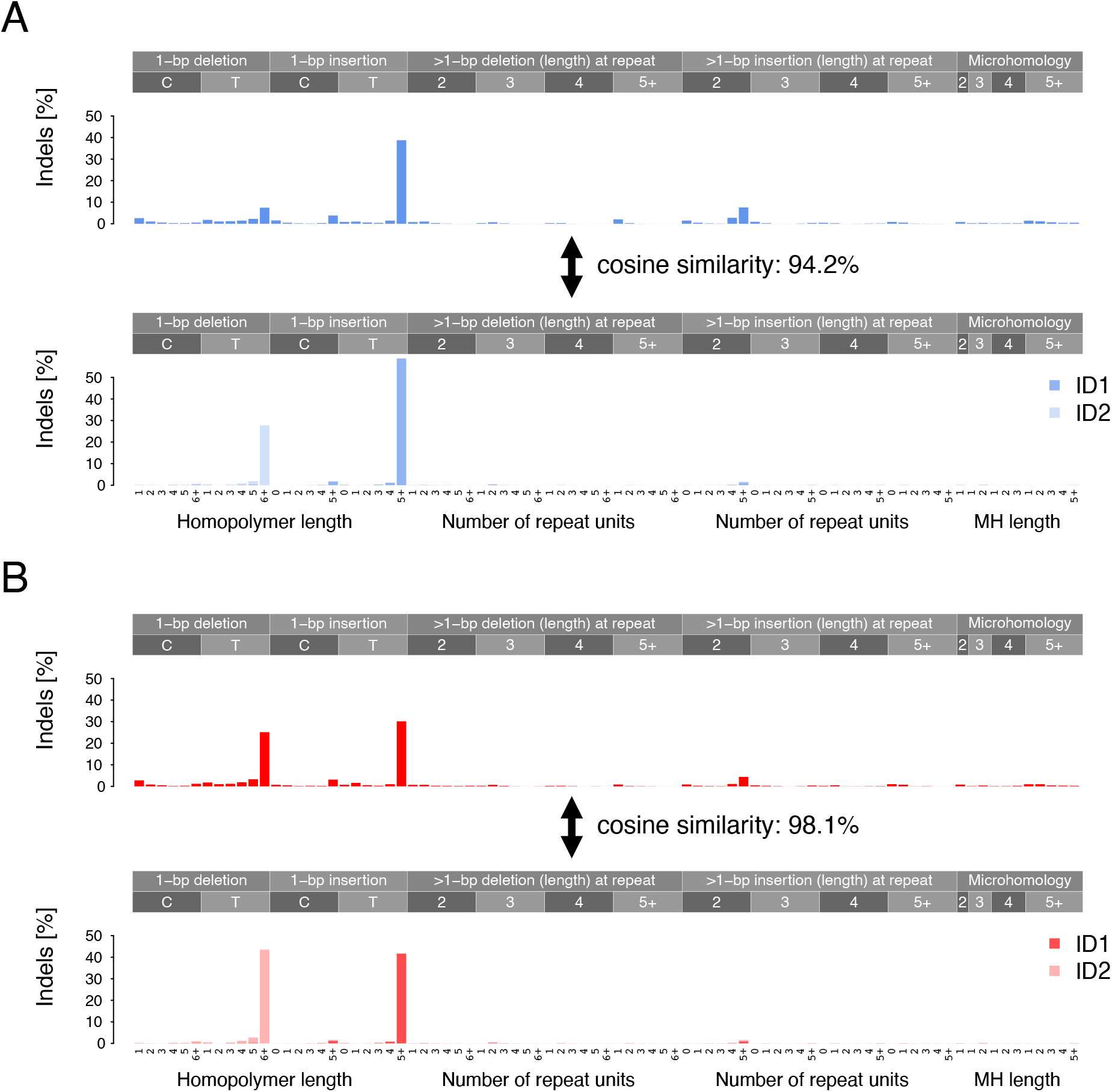
Indel mutational spectra in DFT1 and DFT2. High resolution labelled version of DFT1 (**A**) and DFT2 (**B**) plots presented in Figure 3B. Cosine similarities with spectra reconstructed with COSMIC signatures ID1 and ID2 are shown.

**Fig. S9.**
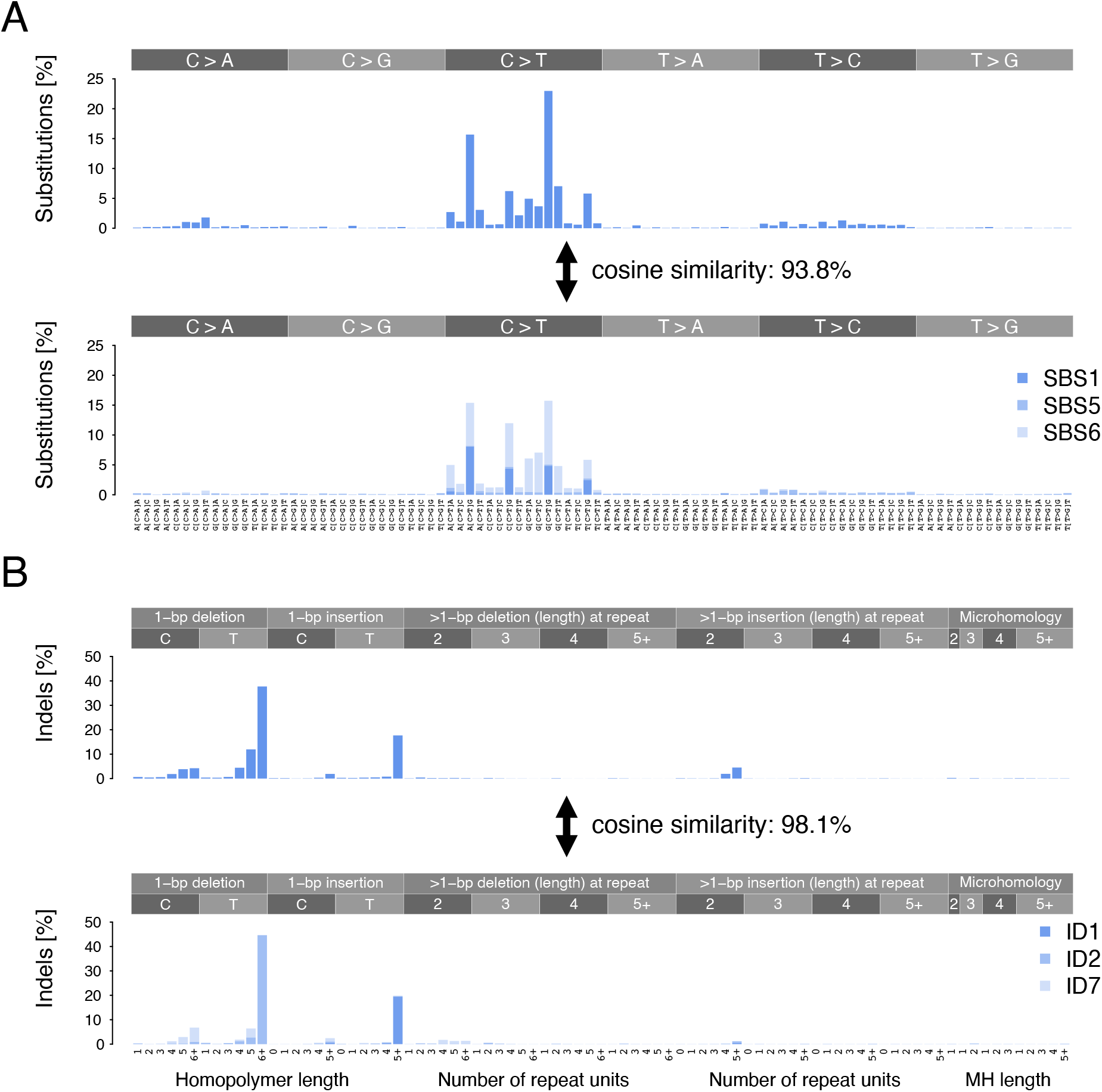
Substitution and indel mutational spectra of 377T1. High resolution labelled version of 377T1 substitution (**A**) and indel (**B**) mutational spectra presented **in Figure 3H.** Cosine similarities with spectra reconstructed with COSMIC signatures SBS1, SBS5 and SBS6 (A) and ID1, ID2 and ID7 (B) are shown.

**Fig. S10.**
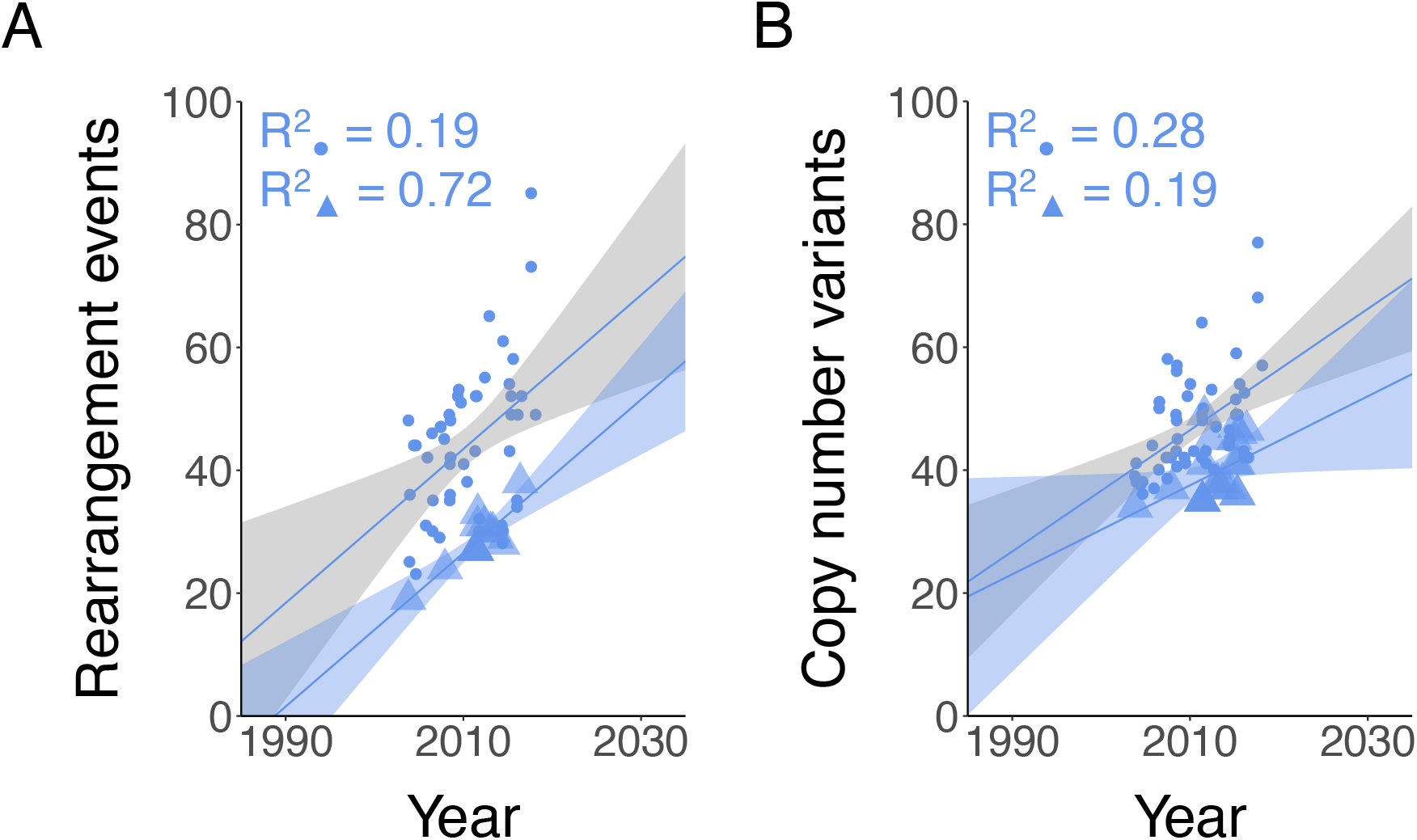
Rearrangement event and copy number variant mutation burden in DFT1 clade C2/3 tumours. Number of rearrangement events (**A**) and copy number variants (**B**) are plotted by sampling date for DFT1 tumours. Tumours belonging clade C2/3 are represented with triangles. All other tumours are represented with circles. Linear regression lines are shown, together with 95% confidence intervals.

**Fig. S11.**
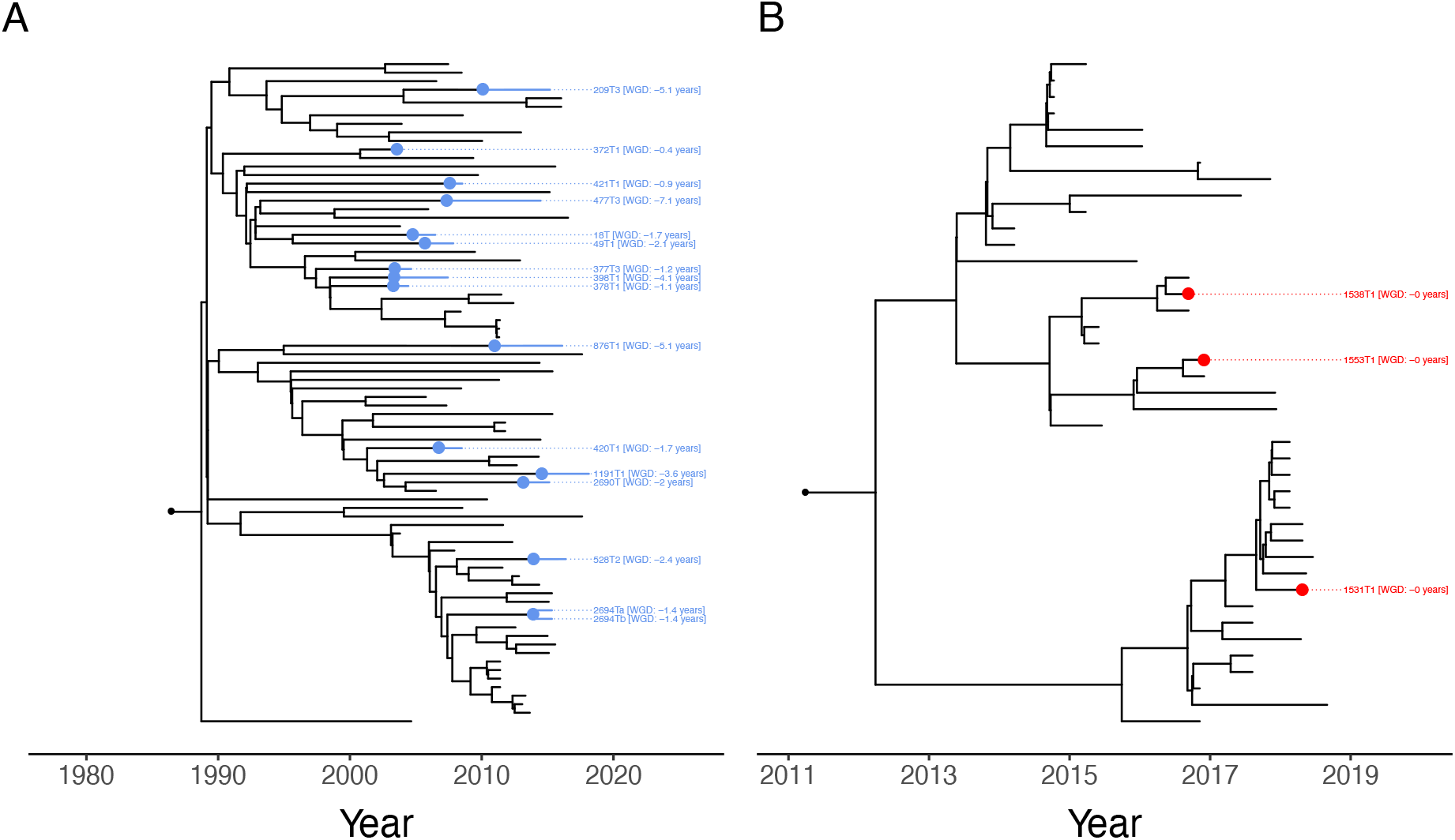
Tetraploidy in DFT1 and DFT2. High resolution labelled version of trees presented in Figure 5F for DFT1 (**A**) and DFT2 (**B**). Tips corresponding to tetraploid tumours are labelled with sample name and the estimated date of whole genome duplication (WGD) in years before sampling. Dots on tree represent mean estimated date of whole genome duplication.

### 9. Supplementary tables

**Table S1. mSarHar1.11 genome assembly information**

(A) Sequencing metrics for reads generated using Oxford Nanopore Technology (ONT). Coverage refers to sequencing depth aligned to mSarHar1.11. BP, base pairs.

(B) Length of physical sequence captured by ONT, 10x linked-reads and Hi-C sequencing technologies. “Range start” and “Range end” refer to a genomic interval length. The proportion of units for each sequencing technology spanning intervals of the associated length are indicated. Units refer to: read length (ONT), size of a barcoded segment (10x linked-reads) and length spanned between junctions (Hi-C reads). BP, base pairs.

(C) Contig assembly benchmark of six algorithms (*105–109*) for Tasmanian devil nanopore long-read sequences. wtdbg (v1 (*105*)) produced the most contiguous assembly.

(D) Curated reference genome assembly statistics from NCBI RefSeq (mammals) and the Vertebrate Genome Project (VGP).

(E) Coordinates of p-arms, q-arms and centromeres for Tasmanian devil chromosomes in mSarHar1.11. MBP, megabases.

(F) Details of tissue samples used to generate a multi-tissue transcriptome atlas for use in mSarHar1.11 gene annotation. Tissues were collected into RNAlater (Sigma-Aldrich, St. Louis, USA), followed by RNA extraction. RNA integrity values were generated using an Agilent Bioanalyzer (Agilent Techologies, Santa Clara, United States).

(G) Counts of genes belonging to different classes annotated in mSarHar1.11 in Ensembl v105. These are compared with equivalent counts from several comparator genomes (all Ensembl v105, except for DEVIL7.0 which was v101). Note that comparator annotations were generated at different times and using different levels of transcriptomic data.

(H) Counts of transcripts belonging to different gene classes annotated in mSarHar1.11 in Ensembl v105. These are compared with equivalent counts from several comparator genomes (all Ensembl v105, except for DEVIL7.0 which was v101). Note that comparator annotations were generated at different times and using different levels of transcriptomic data.

(I) Counts of protein-coding genes belonging to different classes annotated in mSarHar1.11 (Ensembl v105). These are compared with equivalent counts from several comparator genomes (all Ensembl v105, except for DEVIL7.0 which was v101). ‘Genes in a tree’, protein-coding genes that were placed in a multi-species gene tree. ‘Orphans’, genes that could not be placed in a multi-species gene tree. ‘Split genes’, genes likely split into two separate genes when compared to orthologues in other species, usually due to mis-annotation. ‘Short genes’, genes where the longest translation of the transcripts from the gene appears to be truncated when compared to orthologous sequences. ‘Long genes’, genes where the longest translation appears to be expanded when compared to orthologous sequences.

(J) Coordinates of full-length (>6.3 kb) LINE-1 elements in mSarHar1.11. Annotation was performed using RepeatMasker v4.0.8 (*96*).

**Table S2. Sample metadata.**

Details of tumour and normal Tasmanian devil genomes analysed in this study. For cell lines, “date of sampling” refers to date upon which cell line was established. Tumour purity estimates determined using methods involving copy number variants (CNV) and somatic variant allele fraction (VAF) are shown. A pair of DFT1 samples (837T1a, 2695T) from the same individual (microchip ID 982000123160259), supplied by (*110*), were reported to have been collected in distant locations (Takone and Wilmot, respectively) within three months. Based on our phylogenetic tree (Figure S3), we find it probable that both tumours were sampled from a single animal in Takone. Metadata for 2695T have therefore been marked with an asterisk., and the animal ID 837 added in parentheses.

**Table S3. Substitution and indel variant data.**

(A) Details of numbers of substitution and indel variants removed during filtering steps. “DFT1 somatic – shared ancestral” and “DFT2 somatic – shared ancestral” refer to variants shared among all tumours within the DFT1 or DFT2 lineage, respectively. Full lists of genotyped DFT1 somatic, DFT2 somatic, 340T somatic, as well as germline variants after copy number polymorphism filtering are provided in Supplementary Data S1.

(B) Heterozygosity, expressed as heterozygous sites per kilobase (HET SNPs/KB) in Tasmanian devil tumour and normal samples. Only diploid tumours with matched host available and purity >75% were included in the tumour analysis. The number of heterozygous germline single nucleotide polymorphisms (SNPs) was determined in each genome by counting SNPs falling between custom lower (HOM REF / HET VAF THRESHOLD) and upper (HET / HOM ALT VAF THRESHOLD) variant allele fraction (VAF) thresholds. Only autosomal regions of copy number 2 were considered, and the length of the analysed region for each sample is indicated (HIGH QUALITY CN2 SEGMENTS, BP).

(C) Counts of somatic substitutions in each tumour. Sample-specific lower variant allele fraction (VAF) thresholds used during variant calling are shown. Substitution counts normalised for genome opportunity in tetraploid tumours are shown. Counts of mutations assigned to COSMIC signatures SBS1 and SBS5 are shown, alongside cosine similarity of each tumour’s substitution spectrum reconstructed with two subsets of substitution mutational signatures (SBS1, SBS5) and (SBS1, SBS5, SBS6).

(D) Counts of somatic indels in each tumour. Sample-specific lower variant allele fraction (VAF) thresholds used during variant calling are shown. Indel counts normalised for genome opportunity in tetraploid tumours are shown. Counts of indel mutations assigned to COSMIC signatures ID1 and ID2 are shown, alongside cosine similarity of each tumour’s indel mutational spectrum reconstructed with two subsets of mutational signatures (ID1, ID2) and (ID1, ID2, ID7).

**Table S4. Somatic LINE-1 insertions in DFT1 and DFT2.**

(A) Counts of somatic LINE-1 insertions in Tasmanian devil tumours. Counts of two tumours, as indicated by asterisks, only reflect hits identified with SvABA and/or MSG, but not RetroSeq.

(B) Summary of DFT1 somatic LINE-1 insertions. LINE-1 insertions were predicted with RetroSeq, SvABA and/or MSG. LINE-1 insertions that are not predicted to carry 3’ transductions are denoted “solo-L1”. “Original POS1” and “Original POS2” refer to breakpoint coordinates before TIGRA-SV reassembly. “TIGRA POS1” and “TIGRA POS2” refer to breakpoint coordinates after TIGRA-SV reassembly. Genes with exons (“-En”, with n referring to the ID of the exon involved) or introns annotated at a predicted insertion site are indicated; “NCBI” refers to RefSeq v103, “ENSEMBL” refers to Ensembl v104. Those genes contained within the COSMIC Cancer Gene Census version v94 are noted in the column labelled “COSMIC”. “QUAL”, “FILTER” and “INFO” refer to the original VCF entries outputted by Manta, SvABA or RetroSeq. Breakpoint assembly performed using Manta and/or TIGRA-SV are listed, where successful. Genotypes are noted in each DFT1 tumour in columns labelled with each tumours’ ID. The format of the genotype differs among callers: insertions called only with RetroSeq are blank in the case of absence, or display the number of reads supporting the insertion in the case of presence; insertions called with SvABA display the number of reads supporting the insertion; insertions called with Manta display the number of reads supporting the insertion as a fraction of total reads covering the site.

(C) Summary of DFT2 somatic LINE-1 insertions. Notation is as described in (B).

(D) Summary of somatic LINE-1 insertions occurring in 340T, the non-transmissible anal sac carcinoma. Notation is as described in (B).

**Table S5. Somatic rearrangements in DFT1 and DFT2.**

(A) Counts of somatic rearrangements in Tasmanian devil tumours.

(B) Summary of DFT1 somatic rearrangements. Rearrangements were predicted with SvABA and/or MSG. Groups of clustered and phylogenetically concordant rearrangements were collapsed into “rearrangement events”. Each rearrangement has two predicted breakpoints, with coordinates denoted “POS1” and “POS2”, respectively. Breakpoint reassembly was performed using TIGRA-SV, yielding “TIGRA POS1” and “TIGRA POS2” coordinates and a TIGRA-SV assembly breakpoint type (microhomology, blunt end fusion, non-templated sequence insertion). TIGRA-SV breakpoint reassembly sequence is shown, along with alternatives if applicable. Genes annotated by NCBI (RefSeq v103) or ENSEMBL (Ensembl v104) whose exons or introns overlapped predicted breakpoints are noted. The identity of the exon or intron involved, as well as the phases (“geneframe”) of the exons predicted to precede or follow the breakpoint are shown. The copy number of each breakpoint are shown; if there is variation in copy number states among tumours, then tumours with each copy number state are indicated in parentheses. “QUAL”, “FILTER” and “INFO” refer to the original VCF entries outputted by Manta or SvABA. Breakpoint assembly performed using Manta and TIGRA-SV are listed, where successful. Genotypes are noted in each DFT1 tumour in columns labelled with each tumour’s ID. The format of the genotype differs among callers: rearrangements called with SvABA display the number of supporting reads; rearrangements called with Manta display the number of reads supporting the rearrangement as a fraction of total reads covering the site. For each SV, the breakpoint entry with highest “QUAL” or highest overall read support is displayed.

(C) Summary of DFT2 somatic rearrangements. Notation is as described in (B).

(D) Summary of somatic rearrangements occurring in 340T, the non-transmissible anal sac carcinoma. Notation is as described in (B).

**Table S6. Somatic copy number variants in DFT1 and DFT2.**

(A) Counts of somatic copy number variants (CNVs) in Tasmanian devil tumours. The CNV count in the non-transmissible carcinoma, 340T, has not been curated to account for multiple step changes and is marked with an astersisk.

(B) DFT1 somatic copy number variants (CNVs). Each CNV is annotated with an ID. Independent events involving the same genomic interval (for example due to recurrence or backmutation or step changes >1) have separate IDs. Each CNV is associated with a genomic interval, and is annotated as a loss or gain. CNVs unique to tetraploid tumours are noted as “pre-tetraploidisation” or “post-tetraploidisation”. N indicates the number of tumours carrying the CNV, and genes annotated within CNV interval by NCBI RefSeq v103 or Ensembl v104 are displayed. Genes annotated in the COSMIC Cancer Gene Census v94 are listed in the “COSMIC” column. CNVs associated with marker 5 (*18*) or chromosome 2 double minutes are noted. Genotypes are noted in each DFT1 tumour in columns labelled with each tumours’ ID with presence represented with “1” and absence by “0”.

(C) DFT2 somatic CNVs. Notation is as described in (B).

(D) 340T somatic CNVs. Notation is as described in (B), but without splitting multi-step CNVs into separate events.

(E) Estimated timing of whole genome duplication events in DFT1 and DFT2. Tetraploid tumours are listed in rows. Substitution mutations in DFT1 were inferred to have occurred after tetraploidisation if their VAF was higher than the sample-specific lower VAF threshold (Table S3C) but lower than the “Late CN4 max. VAF threshold” (in the case of copy number 4) or lower than the “Late CN6 max. VAF threshold” (in the case of copy number 6). The “assessed genome” represents the number of base pairs (BP) of autosomal genome found at copy number 4 or 6 in each tetraploid tumour, excluding genome regions masked due to an imbalanced minor copy-number state (i.e. chromosome 6 in 209T3). The number of substitution mutations inferred to have occurred after each whole genome duplication (WGD) event is shown, together with the inferred tetraploidisation date and 95% Bayesian credible intervals. In DFT1, the latter were calculated by applying the BEAST substitution mutation rate (Table 2) to the number of post-WGD mutations, and subtracting the resulting time-interval from the sampling date. In DFT2, no post-tetraploid substitutions were observed, and tetraploidisation date was inferred to be equal to sampling date.

(F) Data underlying the analysis of association between tetraploidy and aneuploidy. Tumours were defined as “aneuploid” if the carried one or more whole-chromosome or whole-chromosome-arm CNVs. CNV IDs of relevant CNVs are listed (see Tables S6B and C). DFT1 cell lines are known to undergo chromosome-arm level CNVs during cell line establishment (*18*), and cell lines were excluded from the analysis. Tumours that were identified as tetraploid purely using copy number information are marked; the reported association between aneuploidy and tetraploidy, identified with a Fisher’s exact test, was detectable even if these tumours were excluded.

(G) Phasing of DFT1 genome regions at seven genomic loci (each denoted by a “CNV phasing interval” ID) observed to be repeatedly involved in CNVs in DFT1. Informative germline SNPs were used to identify the haplotype (“A” or “B”) which was lost or gained in any given event. CNVs associated with marker 5 (*18*) were not included in this analysis. CNV phasing interval 1 refers to a region of recurrent loss on chromosome 1; the analysis confirms that these losses are not haplotype specific. CNV phasing interval 2 refers to the region of chromosome 2 encompassing *PDGFRB*; this analysis confirms that both parental haplotypes have been amplified in this region. CNV phasing interval 3 refers to a region of recurrent loss on chromosome 3; the analysis confirms that these losses are not haplotype specific. CNV phasing intervals 4 and 5 refer to events associated with whole arm gains and losses on chromosomes 4 and 5 which are predominantly observed in DFT1 cell lines and were previously confirmed to be non-haplotype specific (*18*). CNV phasing interval 6 events refer to repeated gains and losses of a previously reported (*18*) 12.7 Mb region of chromosome 5. Our analysis identifies this event to be haplotype-specific, and suggest that rearrangements occurring early in DFT1 evolution embedded a single haplotype of this chromosome 5 interval into a repetitive region. It appears that further rearrangements involving flanking repeats have occasionally caused the rearranged haplotype to undergo copy number gains and losses. CNV phasing interval 7 events refer to a series of repeated gains involving a small region of chromosome X. The analysis reveals these to be haplotype-specific. The mechanism in this case remains unexplained.

**Table S7. Annotation of somatic substitutions and indels in DFT1 and DFT2.**

(A) Ensembl Variant Effect Predictor (*88*) output for DFT1 somatic substitutions and indels, annotated relative to Ensembl v104 and RefSeq v103 (only Ensembl details are shown when the same gene is represented in both annotations). COSMIC Cancer Gene Census v94 hits are flagged in the “COSMIC” column. Genotypes are noted in each DFT1 tumour in columns labelled with each tumour’s ID, represented by the number of reads supporting the mutation as a fraction of total reads covering the site.

(B) Substitution and indel mutation gene annotation in DFT2. Notation is as described in (A).

(C) Substitution and indel mutation gene annotation in 340T, the non-transmissible anal sac carcinoma. Notation is as described in (A).

**Table S8. dNdS in Tasmanian devil tumours.**

(A) dNdS summary of genic variants in DFT1, DFT2 and 340T, the non-transmissible anal sac carcinoma, as annotated by *dNdScv* (*33*). “MLE” column denotes maximum likelihood estimates of dNdS for each lineage, and “CI low” and “CI high” indicate the lower and upper 95% confidence intervals, respectively.

(B) Gene-level dNdS summary of DFT1 mutations, output from *dNdScv* (*33*). n_syn, n_mis, n_non, n_spl and n_ind list the number of synonymous, missense, nonsense, essential splice site and truncating indel mutations involving each gene, respectively. qglobal_cv represents q value from a global likelihood ratio test.

(C) Gene-level dNdS summary of DFT2 mutations, output from *dNdScv* (*33*), details as in (B).

(D) Gene-level dNdS summary of 340T mutations, output from *dNdScv* (*33*), details as in (B).

### 10. Supplementary data sets

#### Data S1. Substitution and indel variant lists

(A) List of somatic substitution and indel variants in DFT1. Genotypes in each tumour are displayed as the number of reads supporting the mutation as a fraction of total reads covering the site.

(B) List of somatic substitution and indel variants in DFT2. Notation is as described in (A).

(C) List of somatic substitution and indel variants in 340T, the non-transmissible carcinoma. Notation is as described in (A).

(D) List of germline substitutions and indels. List excludes variants falling into region of germline copy number polymorphism (Data S2). Notation is as described in (A).

#### Data S2. Germline copy number polymorphisms

Genomic intervals with evidence of germline copy number polymorphism in the Tasmanian devil population. Bins occurring within these intervals were excluded from tumour copy number segmentation.

#### Data S3. Copy number plots

Allele-specific copy number profiles for DFT1 and DFT2 tumours. Total copy number is represented in green, minor copy number is represented in blue.

#### Data S4. Circos plots

Summary plots of absolute copy number profiles for DFT1 and DFT2 tumours, integrated with their respective structural variant genotypes. Total copy number profile is shown as a circular ring track, whereas rearrangements are displayed as arcs connecting chromosome regions in the centre.

### 11. Source data and code deposition

mSarHar1.11 reference genome: https://www.ebi.ac.uk/ena/browser/view/GCA_902635505.1

Sequencing data used in mSarHar1.11 assembly: https://www.ebi.ac.uk/ena/browser/view/PRJEB34649

Ensembl mSarHar1.11 gene annotation: http://www.ensembl.org/Sarcophilus_harrisii/Info/Annotation

RefSeq mSarHar1.11 gene annotation: https://www.ncbi.nlm.nih.gov/genome/annotation_euk/Sarcophilus_harrisii/103/

Sequencing data for multi-tissue transcriptome atlas used in mSarHar1.11 gene annotation: https://www.ebi.ac.uk/ena/browser/view/PRJEB34650

Tumour and normal whole genome sequencing data: https://www.ebi.ac.uk/ena/browser/view/PRJEB51704

Code and functionalities used to process mutation data sets: https://github.com/MaximilianStammnitz/Stammnitz2022

https://github.com/MaximilianStammnitz/SubstitutionSafari

https://github.com/MaximilianStammnitz/Indelwald

https://github.com/MaximilianStammnitz/MSG

